# Sparse graphical modeling for electrophysiological phase-based connectivity using circular statistics

**DOI:** 10.1101/2025.03.04.641567

**Authors:** Issey Sukeda, Takeru Matsuda

**Affiliations:** The University of Tokyo; RIKEN Center for Brain Science

**Keywords:** Circular statistics, Connectivity, EEG, Torus graph, Phase

## Abstract

Identifying phase coupling from electrophysiological signals recorded by multiple electrodes, such as electroencephalogram (EEG) and electrocorticography (ECoG), helps neuroscientists and clinicians understand the underlying brain structures or mechanisms. From a statistical perspective, these signals are multi-dimensional circular measurements that are correlated with one another and can be effectively modeled using a torus graph model designed for circular random variables. Using the torus graph model avoids the issue of detecting spurious correlations. However, the naive estimation of this model tends to lead to a dense network structure, which is difficult to interpret. Therefore, to enhance the interpretability of the brain network structure, this paper proposes a sparse estimation method for the torus graph model using regularized score matching combined with information criteria. In numerical simulations, our method successfully recovered the true dependence structure from a synthetic dataset. Furthermore, we present analyses of two real datasets, one involving human EEG and the other marmoset ECoG, demonstrating that our method can be widely applied to phase-coupling analysis across different types of neural data. Using our proposed method, the modularity of the estimated network structure revealed more resolved brain structures and demonstrated differences in trends among individuals.

## 1 Introduction

Electrophysiological recordings such as electroencephalography (EEG) and electrocorticography (ECoG) provide signals that reflect neural activity with millisecond-level temporal resolution. These techniques have been widely used in both cognitive and clinical neuroscience to study the rapid dynamics of brain function, including sensory processing (e.g., Başar (1988)), attention (e.g., Liu et al. (2013)), and electrical or pathological activities in conditions such as epilepsy (e.g., Leocani and Comi (1999); Smith (2005)). Compared to hemodynamic-based imaging methods like fMRI, EEG and ECoG enable the investigation of fast, transient neural interactions across spatially distributed brain areas. Among the various features that can be extracted from these recordings, oscillatory activity and its temporal properties have been shown to play a critical role in neural computation and communication (e.g., Herrmann et al. (2016)). In particular, increasing attention has been directed toward phase relationships between brain regions, as they are believed to mediate functional connectivity (e.g., Bettus et al. (2008); Khaleghi et al. (2024)) and information transfer (Huang et al., 2015).

Phase synchronization and phase locking — collectively referred to as phase-based connectivity — have been widely studied, as they are thought to underlie communication between neural networks (Singer et al., 1993; Fries, 2005, 2009; Salinas and Sejnowski, 2001; Cohen, 2014). Thatcher et al. (2005) examined the relationship between EEG measures and intelligence, and found that high IQ was associated with shorter frontal phase delays, longer posterior phase delays, and lower coherence — making these connectivity-related features key discriminators between high and low IQ individuals. Busch et al. (2009) investigated the relationship between the phase oscillations and our visual perceptions, showing that oscillatory phase at stimulus onset can predict visual detection performance. Moreover, they have also become important from the perspective of research on brain diseases. For example, the early gamma band phase locking in schizophrenia (Roach and Mathalon, 2008) and the phase synchronization of epilepsy participants (Mormann et al., 2000) have been reported. In addition, various metrics have been developed to quantitatively measure phase relationships, such as the phase locking value (PLV) (Lachaux et al., 1999) and the phase lag index (PLI) (Stam et al., 2007). However, many of the these approaches are relatively simplistic or descriptive statistical methods, which often fail to account for the full complexity of phase relationships.

From a statistical perspective, phase data are inherently circular, as they lie on the unit circle. To handle such circular data effectively, specialized statistical methods are required. Here, we introduce circular statistics, a branch of statistics specifically designed to analyze angular or circular data, to offer a robust framework for analyzing phase relationships (e.g., Mardia (1975); Mardia and Jupp (2009); Ley and Verdebout (2018)). Circular statistics involves the application of circular distributions such as the von Mises distribution, a circular analogue of the normal distribution, and techniques such as circular correlation and angular regression, which are well-suited to analyzing angular data. Traditionally, these methods have been widely applied in fields such as meteorology and environmental sciences, for example in the analysis of wind direction(e.g., Carta et al. (2008), Masseran et al. (2013)). Although these methods are suitable for analyzing angular data, their application in electrophysiological phase analysis in neuroscience remains limited. One notable exception is Klein et al. (2020), who applied the torus graph, a high-dimensional circular model, to analyze local field potentials collected from the prefrontal cortex and hippocampus of a macaque monkey.

This study applies circular statistics to enhance the analysis of phase dependencies in EEG data, addressing their underutilization in this context. Specifically, we apply a graphical modeling approach, which is a statistical tool that use graphs to represent conditional independence among variables; nodes correspond to variables, and edges indicate dependencies. We adopt this approach by statistically estimating the torus graph (Klein et al., 2020), a graphical model designed to capture conditional dependencies among circular variables, such as phase angles derived from EEG/ECoG recordings. Our proposed method, summarized in Fig. 1, simultaneously handles multiple phase signals and naturally yields a sparse, interpretable graph network structure without relying on arbitrary binarization or thresholding — common practices that often compromise result reliability in neuroscience. The resulting graph network can be interpreted as the brain’s functional connectivity. To analyze these structures, we examined key graph properties such as modularity and small-worldness. Our method is applied to both synthetic data and real-world electrophysiological data, including 61-dimensional human EEG and 96-dimensional marmoset ECoG, with a focus on modularity differences across participant groups and frequency bands. By utilizing circular statistical techniques, we hope to offer deeper insights into the phase dynamics of brain waves and their potential implications for understanding neural communication and brain function.

**Figure 1:**
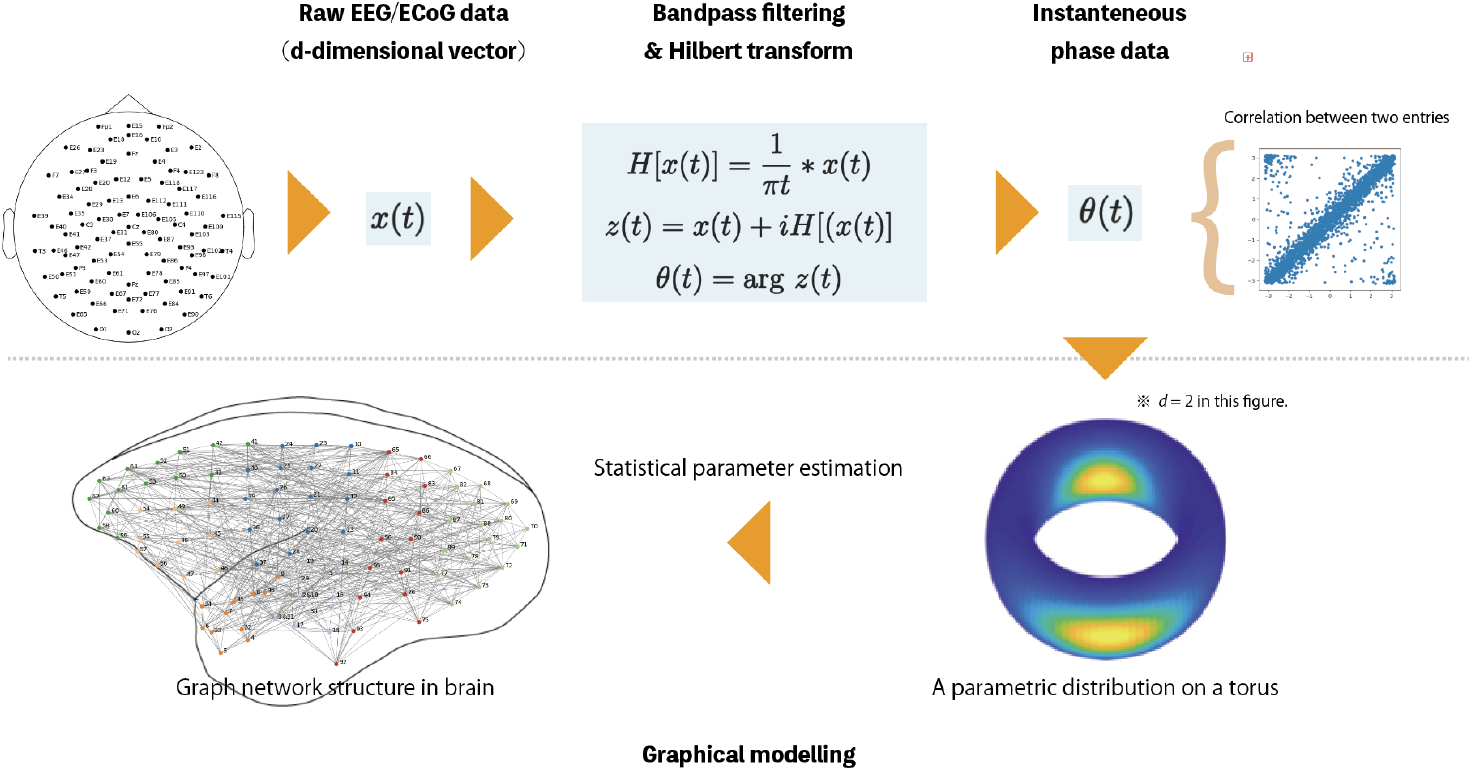
Overview of graphical modeling of EEG/ECoG phase data using a distribution on a torus.

## 2 Method

### 2.1 Data acquisition

The data used in our paper are all publicly available datasets from existing studies, and we have not collected any new data independently. Firstly, we used human EEG data with 91 measurement channels published by Chennu et al. (2016) and extracted 61 channels with a length of 25000 (5 seconds at 500Hz, repeated for 10 epochs) for analysis. According to Chennu et al. (2016), these data were collected from 17 participants. Among those participants, seven were labeled as a “drowsy” group and the rest as a “responsive” group, depending on the perceptual hit rate after administration of propofol. Each participant’s EEG was recorded during the following four states: baseline, mild, moderate, and recovery. We also used two marmoset ECoG datasets provided by Komatsu and Ichinohe (2020, 2021) with all 96 measurement channels. The marmoset IDs are R03 0024 CM622M and R03_ 0035_ CM814F. The types of conditions in these data were as follows: AD-Awake, AD-K10, AD-K30, and AD-K150, where K10, K30, and K150 mean 10, 30, and 150 minutes after ketamine administrations, respectively. We extracted each data such that its length is 25000 starting from the beginning. According to the original paper by Komatsu and Ichinohe (2021), these data were collected from two marmosets (one from the left hemisphere and another from the right hemisphere of their brains).

### 2.2 Torus graph model for high-dimensional circular data

Recently, Klein et al. (2020) proposed the torus graph model^1^, which is a high-dimensional circular distribution. It is named after the torus, a higher-dimensional generalization of the circle in mathematics and topology.

Suppose *X* is a *d*-dimensional circular random vector. The torus graph model is defined by the following exponential family parametrized by *ϕ*:

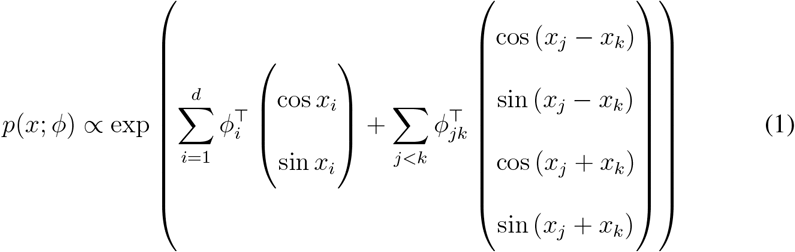

where *ϕ* = [{*ϕ*_*i*_}_*i*_, {*ϕ*_*jk*_}_(*j,k*)_]. We refer to this as the full torus graph model, in contrast to the submodels including the rotational model where *ϕ*_*jk*_ = (0, 0, ·, ·)^⊤^ and the reflectional model where *ϕ*_*jk*_ = (·, ·, 0, 0)^⊤^ (see Fig. 2). This model is not a time-series model but an i.i.d. formulation. Within this framework, the observed samples are pooled across time for estimation, even though EEG/ECoG raw recordings are inherently time-series data. Our subsequent approach inherits this aspect in the same manner.

**Figure 2:**
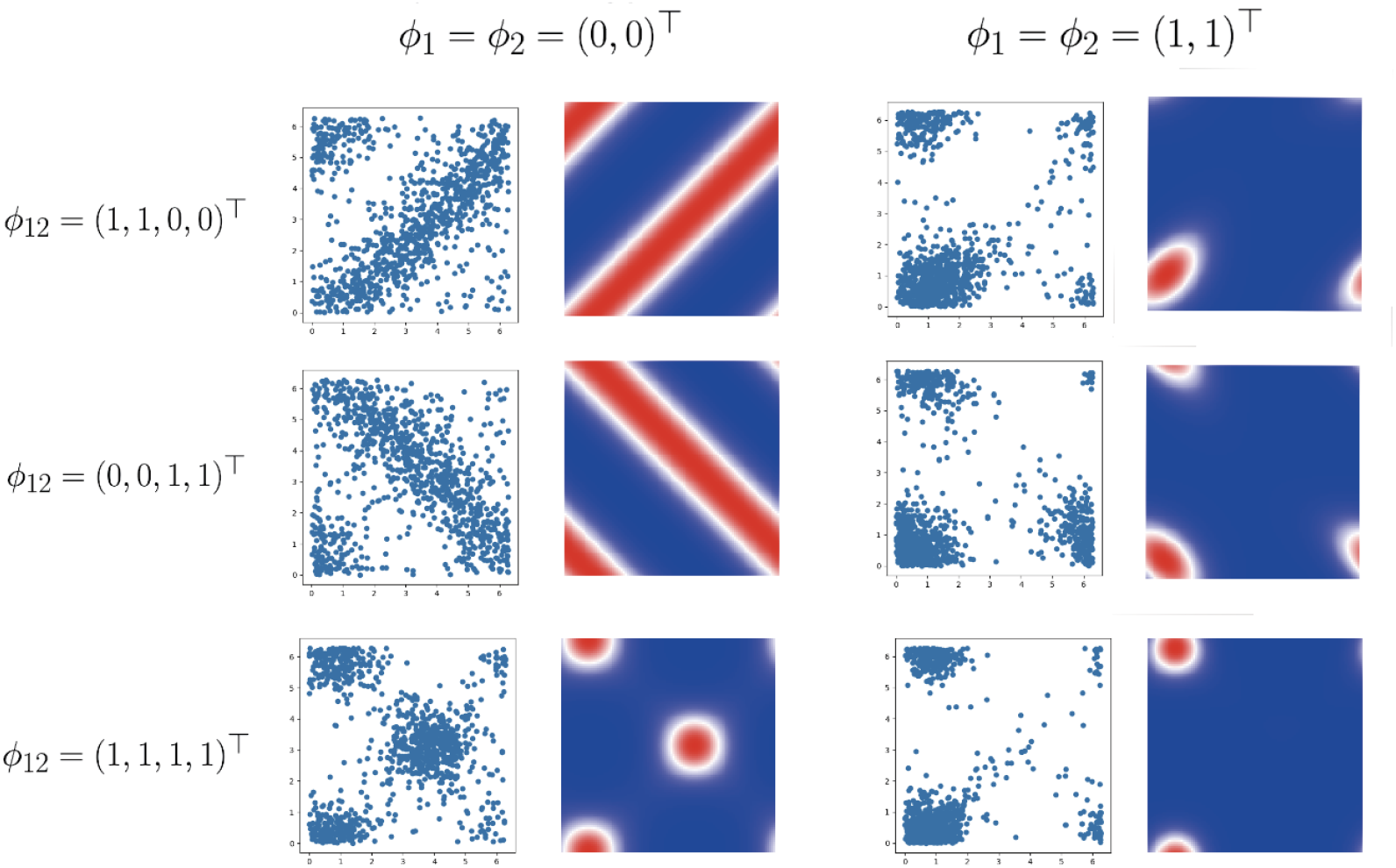
Submodels of torus graph model. 1000 samples and the density function are plotted for each submodel of the torus graph model.

Notably, the torus graph model is a parametric non-normalized model, and its parameters can be easily estimated using score matching since it belongs to an exponential family by its definition (Hyvärinen, 2005). The model incorporates the interactions between different channels through pairwise terms, making it naturally adaptable to graphical modeling. Although the originally proposed torus graph model is quite flexible and applicable when applied to high-dimensional circular data, an estimated result by naive score matching yields a dense graph, which makes it challenging to directly interpret the estimated functional connectivity between different brain regions.

To understand which parts of the brain are relatively active or which connections are stronger than others, it is preferable to obtain a sparse graphical structure. We remark that the assumption that the underlying graph is sparse is not essential but rather adopted for convenience, primarily because dense graphs are often considered uninterpretable especially when visualization is the main scope. In practice, among neuroscience researchers, a common approach to obtaining a sparse graphical structure is to apply a threshold — the connections stronger than the threshold are retained, whereas weaker ones are discarded. However, this method introduces an arbitrary error into the final result, as the outcome is highly sensitive to where the choice of threshold levels are set. Such sensitivity can significantly affect the final result, thereby reducing the reliability of the overall estimation. To address this issue, Klein et al. (2020) performed statistical testing on each edge individually based on the dense estimator to determine which edges should be retained. Bonferroni correction analysis was performed to mitigate the effects of multiple testing. Our method also seeks to remove weaker edges; however, it does so in a principled manner, guided by a combination of regularization and information criteria, rather than arbitrarily.

### 2.3 Regularized score matching method for the torus graph model

While the aforementioned approaches rely on dense estimators, another direct approach is to allow the model to induce a sparse estimator, which includes many zero entries. Regularization is the typical method used to achieve this in machine learning. Here we propose to incorporate the regularization to the torus graph model (Klein et al., 2020).

In the torus graph model, *ϕ*_*jk*_ = (0, 0, 0, 0)^⊤^ is equivalent to the conditional independence of *x*_*j*_ and *x*_*k*_; *x*_*j*_ and *x*_*k*_ are independent given all other variables (Corollary 2.1-2 of Klein et al. (2020)). Hence, this model can be used for graphical modeling. Suppose *G* = *G*(*V, E*), *V* = {1, · · ·, *d*}. The natural parameters in the torus graph model (1) can be interpreted as follows: *ϕ*_*i*_ is a two-dimensional vector corresponding to the node *i*, whereas *ϕ*_*jk*_ is a four-dimensional vector representing the interaction between the node *j* and *k*. Then, the problem of identifying the dependence between different nodes reduces to estimating the parameters *ϕ* statistically. Note that the normalization constant of (1) is intractable, so the usual maximum likelihood estimation is also intractable. In such cases, score matching estimation is known to be useful. Specifically, the score matching objective of the exponential family takes the quadratic form (Hyvärinen, 2005). For the torus graph model, the objective function is known to be

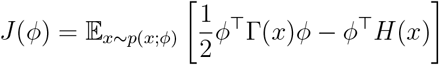

where

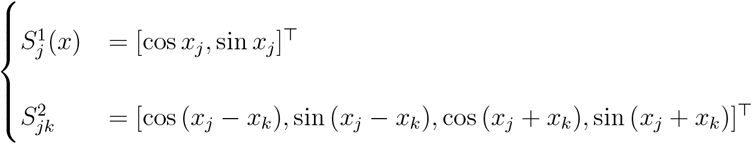

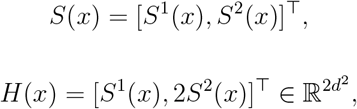

and

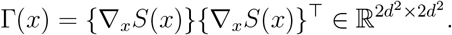

On the basis of above formula, the empirical objective function using *N* observed sam-ples becomes

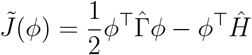

where

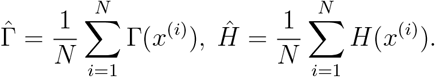

As a result, the score matching estimator of *ϕ*, which is the minimizer of 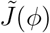, is easily calculated by using the formula below:

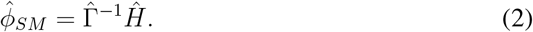

See Klein et al. (2020) for the explicit expressions for 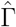 and *Ĥ*.

The approach for determining the graph structure via naive score matching of the torus graph model entails the issue of arbitrariness in binarization. Since the full torus graph (resp. rotational model) assigns four (resp. two) estimated values for each of the *d*(*d* − 1)*/*2 edges, this approach tends to complicate the interpretation of the estimated results. Therefore, the binarization process is conducted to recover the underlying graph structure and make the results easier to interpret. The easiest method is to calculate the norm of estimated vectors corresponding to each edge, and then pull an edge if only the norm surpasses a certain threshold. However, there remains an arbitrary choice of determining an exact threshold value. To avoid this issue, Klein et al. (2020) conducted hypothesis testing that examines the null hypothesis *ϕ*_*jk*_ = 0 for each (*j, k*) to determine the dependence of each edge and recover an interpretable graph structure. Although this approach is statistically acceptable, the number of hypothetical tests grows exponentially under general settings. This issue was not a crucial problem in their work as they used the pre-given community structure of the network from the well-known domain knowledge, however, such prior knowledge is not always available.

To address this issue, *l*_1_-regularization methods can be applied, especially for higher dimensional problems, leading to a sparse solution.^2^ Similar approaches for the general exponential families were studied by Lin et al. (2016). Following these studies, we propose the use of *l*_1_-regularization for estimating the torus graph parameters by the score matching. This estimator is defined as the minimizer of

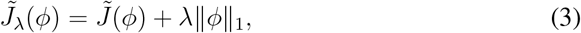

where *λ* is a tuning parameter which controls the sparsity of the resulting solution. Since this objective function being minimized is convex, we may utilize normal optimization methods, such as gradient descent. However, the solution may not remain sparse. Therefore, to address this, a LASSO algorithm (Tibshirani, 1996) such as LARS (Efron et al., 2004) can be implemented. Yet, due to computational efficiency, the alternating direction method of multipliers (ADMM) is a more generally accepted method (Boyd et al., 2011). Our modified objective function, which is required to be minimized with respect to *x*, is

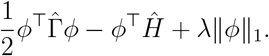

In ADMM, we instead minimize 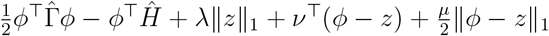, coupled with the dual variable *z* by iterating the following three steps until convergence:

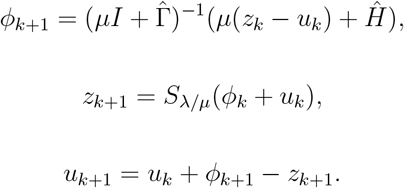

In this process, *I* denotes the identity matrix, *S* denotes the soft threshold function, *z*_*k*_ is the auxiliary parameter, and *µ* is the arbitrary constant.

One of the primary motivations for the use of LASSO algorithms is to obtain a regularization path (or sometimes called the solution path), which is a sequence of solutions along with various *λ* values. Formally, the regularization path is denoted as 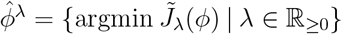, or practically 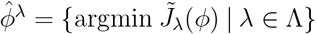 where Λ is the set of candidate values of *λ*. Note that the former becomes piecewise linear, i.e., there exists the change points 0 = *λ*_0_ *< λ*_1_ *<* · · · *< λ*_*R*_ = ∞ and *ξ*_0_, …, *ξ*_*R*−1_ such that 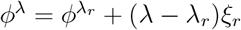 for *λ* ∈ [*λ*_*r*_, *λ*_*r*+1_], but the latter may not.

In practice, we consider the variant problem called Group LASSO (Yuan and Lin, 2006) instead, where the regularization terms are group-wise. In this case, the estimator becomes the minimizer of

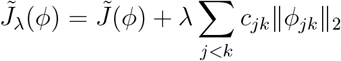

where *c*_*jk*_ is a weight constant assigned to each edge between nodes *j* and *k*. For convenience, we set *c*_*jk*_ = 1 for any corresponding edge (*j, k*). For the optimization process in Group LASSO, the ADMM algorithm requires a slight modification so that the update of *z* becomes group-wise:

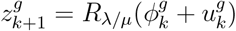

where *g* indicates the partition of parameters into groups, i.e., in our case two dimensional vectors for each node and four dimensional vectors for each edge, and *R* is a shrinkage operator. See Appendix A. for details.

### 2.4 Model selection with score matching information criteria (SMIC)

Although *l*_1_-regularization is a useful method for edge selection in graphical modeling (or variable selection in regression as well), the tuning parameter *λ* is given arbitrarily. To choose an appropriate *λ*, cross validation is widely used but often time consuming, especially when using leave-one-out cross validation. An alternative method is to consider the information criteria, such as the Akaike information criteria (AIC) for the maximum likelihood estimator, and select the model with the smallest AIC. ^3^

In our case, the score matching information criteria (SMIC) proposed by Matsuda et al. (2021) can be used to decide the regularization parameter *λ*, or equivalently the appropriate graph structure. The torus graph model belongs to an exponential family, hence the SMIC is calculated as

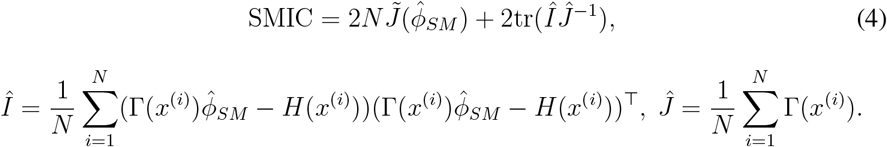

Briefly, the first term 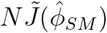 has the negative bias of tr(*I*(*ϕ*)*J*(*ϕ*)^−1^) as the estimator of the predictive distribution *J*(*ϕ*); thus, the bias correction is conducted by adding the second term. To perform model selection via SMIC minimization, we should calculate the value of SMIC for all candidate models and finally select the one with minimum SMIC value. However, since *λ* can vary continuously, we discretize it and select a limited number of points in practice. To effectively identify the best *λ*, we first pick 10 equally spaced discrete points and select an interval that includes the minimum SMIC value. We then repeat the same calculation for this interval recursively. In each recursive iteration, parallel computing can be applied since the calculation of SMIC is independent for each choice of *λ*, which further accelerates the entire procedure.

### 2.5 Graphical properties of the estimated network structure

Once the appropriate graphical model is specified, the representative graphical properties can be observed. Indeed, we obtained the following four metrics: (1) number of edges, (2) average clustering coefficient, (3) average shortest path length, and (4) modularity.

Average clustering (Schank and Wagner, 2005) measures how often the neighbors of a node are connected to each other. For an undirected, unweighted graph *G* = (*V, E*), let *m*(*v*) = |{(*u, w*) ∈ *E* | (*v, w*) ∈ *E* and (*v, u*) ∈ *E*}| and 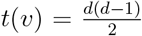 where *d* is the degree of node *v*. Then, the average clustering coefficient is defined as

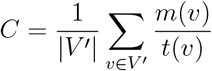

where *V* ^*′*^ is the set of nodes *v* with *d* ≥ 2. As the value of *C* (∈ [0, 1]) becomes larger, the more densely connected the network becomes.

Average shortest path length measures the length of paths between any two nodes in the network. For a connected undirected unweighted graph *G* = (*V, E*),

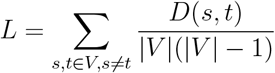

where *D*(*s, t*) denotes the shortest path length between node *s* and *t*. The smaller *L* (≥ 1) is, the more densely connected the network is.

In addition, community-detection-based measures are also considered useful. Newman and Girvan (Newman and Girvan, 2004) introduced the modularity maximization problem for community detection. For an undirected, unweighted graph *G* = (*V, E*), |*E*| = *m*, modularity is defined as

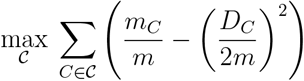

where 𝒞 is a clustering (a set of mutually exclusive sets of *V*), *C* denotes each cluster, *m*_*C*_ denotes the number of edges in *C, D*_*C*_ denotes the sum of degree of nodes in *C*. The larger the modularity is, the denser the connections within each cluster become, and the sparser the connections become between the different clusters in the network. To overcome the drawbacks of the modularity (Fortunato and Barthelemy, 2007; Good et al., 2010), Li et al. (2008) proposed the modularity density maximization problem. Other groups have tested this method and have developed mathematical approaches to improve the computation burden and heuristic methods (Costa, 2015; Sukeda et al., 2023). Among them, calculating modularity by the Louvain method (Blondel et al., 2008) is mostly useful in practice because it operates in time complexity *O*(|*V* | log |*V* |) (Chennu et al., 2017; Lui et al., 2021); hence, we utilize it in the following experiments.

## 3 Simulation results

In this section, we present simulation studies in which our proposed method is applied to synthetic data generated from a known torus graph model (Sections 3.1–3.2) and the Kuramoto model (Kuramoto, 1975) (Section 3.3).

### 3.1 Synthetic data generated from a torus graph

To validate our proposed method, we performed it on synthetic data generated from a prespecified torus graph model containing 5, 19, and 61 dimensions. The purpose of these experiments is to confirm the obvious. The dimensions were chosen arbitrarily to represent different levels of granularity: a coarse, a moderate, and a finer resolution for analysis. Firstly, we describe the procedure in the five-dimensional case; the same applies to other dimensions. To generate synthetic data samples, we use rejection sampling (e.g., Von Neumann (1951), Robert et al. (2010)) with uniform proposal to generate 1000 data points from a torus graph model: draw *x* uniformly over the support and accept it with probability *p*(*x*; *ϕ*)*/M*, choosing 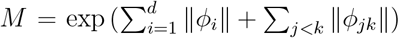. Its true graph structure here is set to *G* = *G*(*V, E*), *V* = {1, 2, 3, 4, 5}, and *E* = {(1, 3), (1, 4), (2, 4), (2, 5), (3, 5)}. Moreover, the true natural parameter *ϕ* of the torus graph model is set such that the corresponding parameters to *E* are all set to 0.3 and 0 otherwise. Then, we applied our regularized score matching with various *λ* values. For each *λ*, we calculate SMIC, which is plotted in Fig. 3. As depicted in Fig. 4, the SMIC takes the minimum value at the model with five edges, which correctly corresponds to *E*. Conversely, calculating the phase locking value of each edge pair overdetects the edge, suffering from false pseudo correlations.

**Figure 3:**
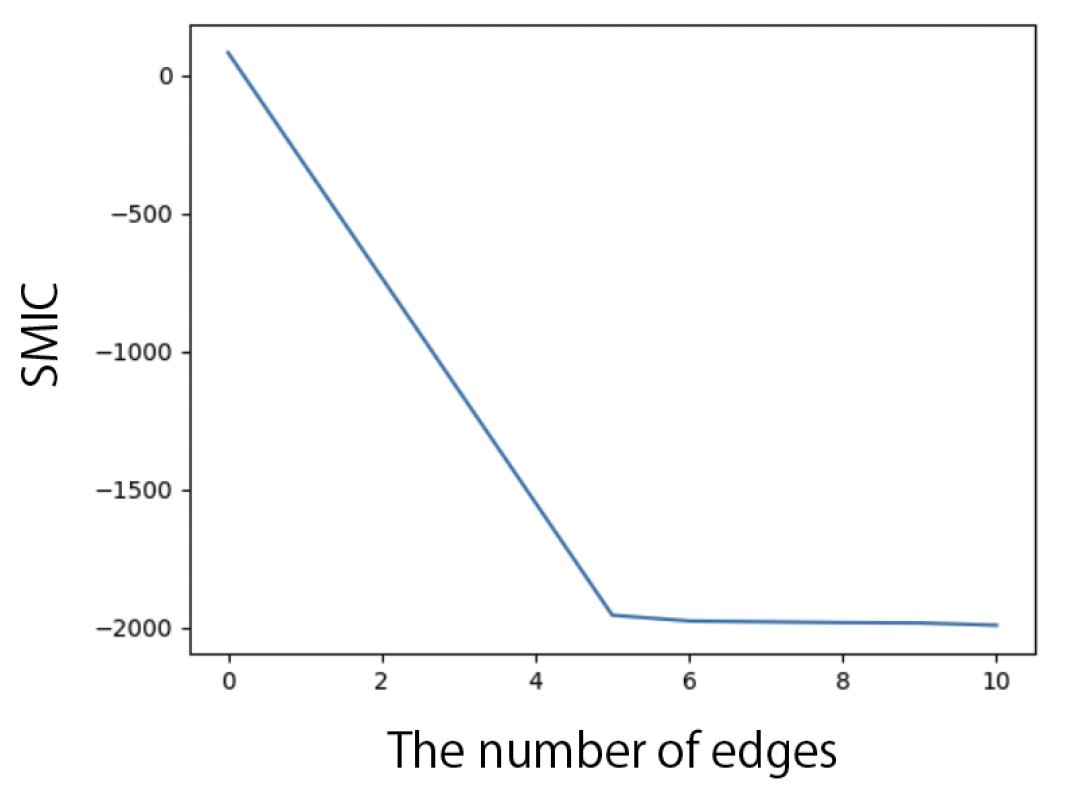
The recovery of graph structure with 5-dimensional synthetic data. Axes are labeled with SMIC (y-axis) and the number of edges (x-axis). The number of synthetic samples is set to 1000 here.

**Figure 4:**
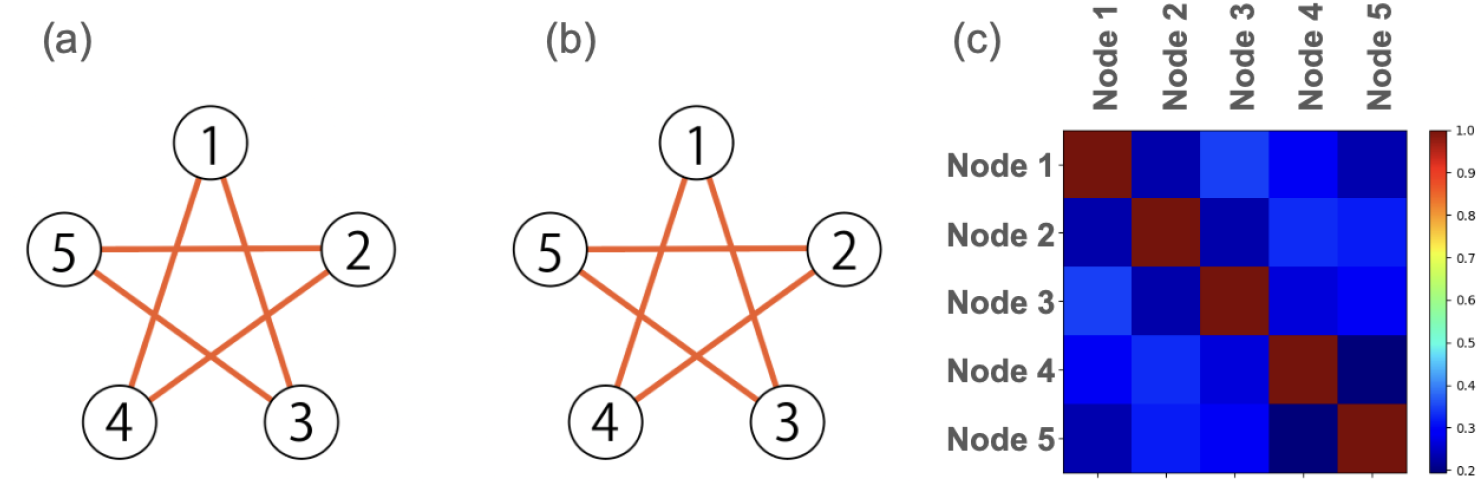
Results on 5 dimensional synthetic data sampled from a predefined torus graph model: (a) the ground truth structure (b) the recovered structure by our method (c) the adjacency matrix representing phase locking value (PLV) between nodes. While the naive phase locking value calculation overdetects the edges, our proposed method correctly recovers the dependence structure.

Next, we generate synthetic data from a high-dimensional torus graph model. Because rejection sampling is computationally prohibitive, we employ Gibbs sampling. Gibbs sampling is a Markov chain Monte Carlo (MCMC) method that draws from a joint distribution by iteratively sampling from its full conditionals. In our model Eq. (1), each full conditional *p*(*x*_*k*_ | *x*_−*k*_; ***ϕ***), *k* = 1, 2, …, *d* is von Mises (see Theorem 2.2 of (Klein et al., 2020) for the explicit derivation), which enables straightforward Gibbs updates. As in standard MCMC practice, we discard an initial burn-in period to allow the chain to approach stationarity.

In this experimental setting, the true model parameter is set as follows. First, we generate a random graph structure named Erdös-Rényi model *G*(*V, E*) = *G*(|*V* |, *p*). In this model, each edge is included in the graph with probability 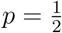. Then we assign

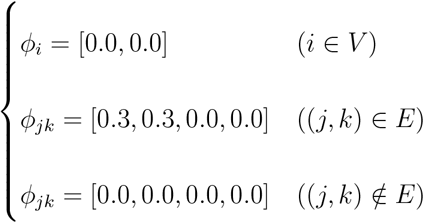

as the ground truth. Note that since this distribution is not multimodal, the simple Gibbs sampling works well. The burn-in period was set to 10000.

As a result, we found that our method based on SMIC minimization tends to select a denser graph structure than the true one. Tab. 1 and Fig. 5 show the results for *d* = 19 and *d* = 61, supported by a permutation test in S4 Appendix. This tendency is consistent with the precedent results, where the model selection methods based on AIC-alike information criteria (Zou et al., 2007) tend to increase the false positive rates. Contrarily, the decline in SMIC drastically changes around the true number of edges: 80 edges in 19 dimensional example and around 889 edges in 61 dimensional example. To avoid arbitrariness, we do not adopt this approach; however, selecting the “elbow” may be useful in practice. See S5 Appendix for simulation-based empirical results.

**Table 1:** Detection probability for high-dimensional synthetic dataset from a random graph structure. Example results for artificially generated dataset sampled from a torus graph where the graph structures are randomly determined by the Erdös-Rényi random graph.

| $d$ | # true edges | # estimated edges | TP | FP | TN | FN | the best $\lambda$ |
| --- | --- | --- | --- | --- | --- | --- | --- |
| 19 | 80 | 122 | 80 | 42 | 49 | 0 | 0.034 |
| 61 | 889 | 1263 | 889 | 374 | 567 | 0 | 1.616 |

**Figure 5:**
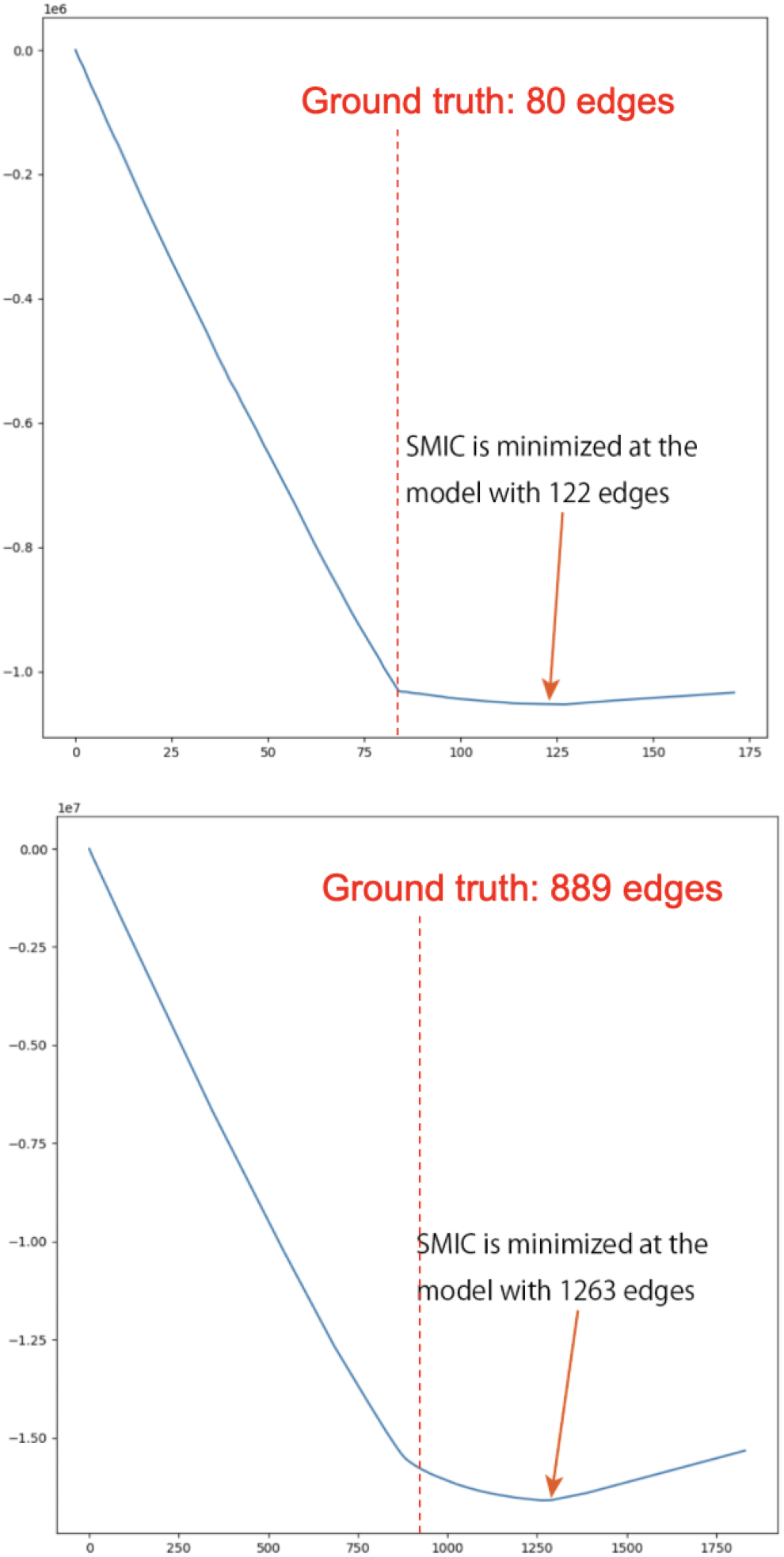
The number of edges vs SMIC for generated dataset sampled from a torus graph. (left: *d* = 19, right: *d* = 61)

### 3.2 Comparison between SMIC and cross validation

We further compare the performance of SMIC minimization against the typical cross validation method to select the regularization parameter *λ*. Here we use the leave-one-out-cross-validation (LOOCV). The counterpart of information criteria, which we call SMCV herein, is calculated as follows:

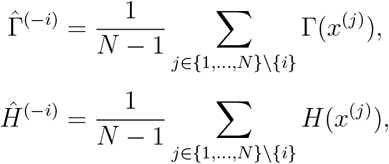

for the SMCV calculation and 0.649 seconds for the SMIC calculation.

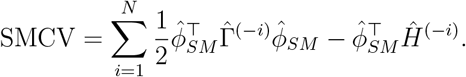

Here, *i* denotes the index of the data, which is left out from the dataset one by one. To compare the performance of SMIC minimization with SMCV minimization, we conducted two simulation experiments using three- and five-dimensional synthetic data, respectively. These simulations show that minimizing SMIC exhibits performance comparable to the standard method of choosing the minimum SMCV model when measured by the edge detection probability, while achieving a significant acceleration as shown in Fig. 6.

**Figure 6:**
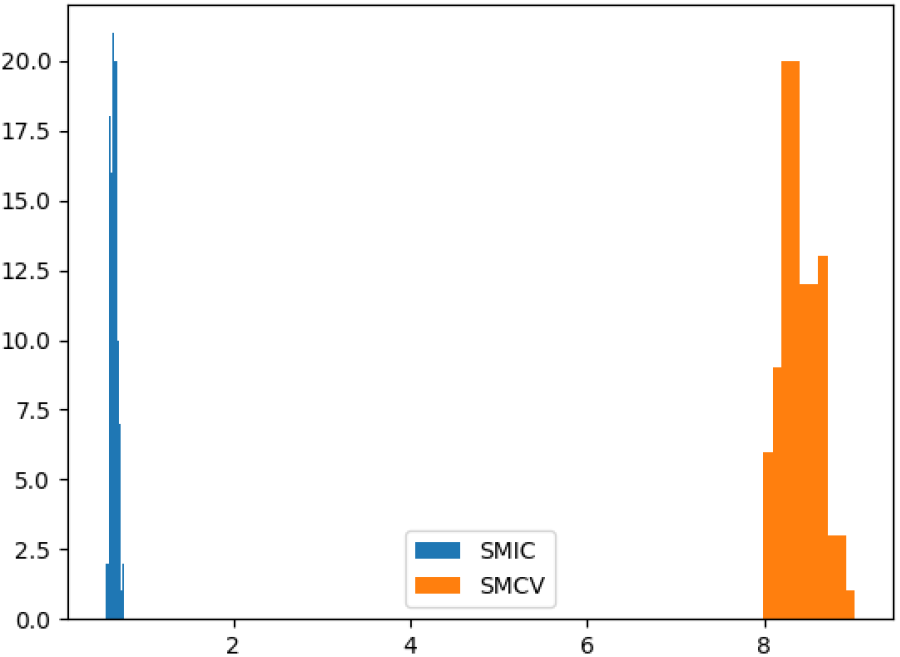
Average computation times in seconds of 30 runs. In average, 8.41 seconds

First, we sample *N* = 100 samples from 3-dimensional torus graph using rejection sampling. Suppose the true graph structure is *G* = *G*(*V, E*), *V* = {1, 2, 3} where *E* = {(1, 2), (2, 3)} and *ϕ* = (0, 0, 0, 0, 0, 0, *c, c, c, c*, 0, 0, 0, 0, *c, c, c, c*)^⊤^, *c* ∈ ℝ. Since *G* consists of 3 edges, the number of candidate models is 2^3^ = 8. We select the model with minimum SMCV or SMIC as the optimal model. Tab. 2 shows the detection probabilities of each edge in the torus graph model for *N* = 100 when *c* = 0.1, 0.2, 0.3, 0.5. As shown in Tab. 3, the similar experiment is also conducted for

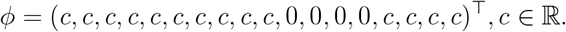

**Table 2:** Detection probability. The 3-dimensional synthetic data were sampled from the torus graph model with *ϕ* = (0, 0, 0, 0, 0, 0, *c, c, c, c*, 0, 0, 0, 0, *c, c, c, c*)^⊤^, *c* ∈ {0.1, 0.2, 0.3, 0.5}.

| $c$ | edge | Ground truth | SMCV | SMIC |
| --- | --- | --- | --- | --- |
| 0.1 | (1,2) | true | 0.426 | 0.430 |
|  | (1,3) | false | 0.201 | 0.203 |
|  | (2,3) | true | 0.436 | 0.437 |
| 0.2 | (1,2) | true | 0.680 | 0.686 |
|  | (1,3) | false | 0.080 | 0.079 |
|  | (2,3) | true | 0.686 | 0.703 |
| 0.3 | (1,2) | true | 0.935 | 0.938 |
|  | (1,3) | false | 0.055 | 0.057 |
|  | (2,3) | true | 0.940 | 0.948 |
| 0.5 | (1,2) | true | 1.000 | 1.000 |
|  | (1,3) | false | 0.043 | 0.078 |
|  | (2,3) | true | 1.000 | 1.000 |

**Table 3:** Detection probability. The 3-dimensional synthetic data were sampled from the torus graph model with *ϕ* = (*c, c, c, c, c, c, c, c, c, c*, 0, 0, 0, 0, *c, c, c, c*)^⊤^, *c* ∈ {0.1, 0.2, 0.3, 0.5}.

| $c$ | edge | Ground truth | SMCV | SMIC |
| --- | --- | --- | --- | --- |
| 0.1 | (1,2) | true | 0.409 | 0.415 |
|  | (1,3) | false | 0.203 | 0.202 |
|  | (2,3) | true | 0.433 | 0.434 |
| 0.2 | (1,2) | true | 0.667 | 0.744 |
|  | (1,3) | false | 0.067 | 0.052 |
|  | (2,3) | true | 0.606 | 0.670 |
| 0.3 | (1,2) | true | 0.843 | 0.950 |
|  | (1,3) | false | 0.041 | 0.060 |
|  | (2,3) | true | 0.834 | 0.952 |
| 0.5 | (1,2) | true | 0.922 | 0.995 |
|  | (1,3) | false | 0.030 | 0.169 |
|  | (2,3) | true | 0.929 | 0.991 |

In both cases, detection probabilities improve monotonically with increasing *c*, which is consistent with the intuition that stronger coupling leads to easier edge detection.

Next, we sample *N* = 100 samples from 5-dimensional torus graph using rejection sampling. Suppose the true graph structure is *G* = *G*(*V, E*),*V* = {1, 2, 3, 4, 5}, *E* = {(1, 3), (1, 4), (2, 4), (2, 5), (3, 5)}. The true natural parameter *ϕ* of the torus graph model is set such that the corresponding parameters to *E* are all set to 0.3 and 0 otherwise, in align with previous experiments. Since the number of full candidate models 2^10^ is too large to explore, we instead employ the group LASSO method to restrict the candidate models into less than 100, which are automatically exploited as the regularization path. The model with least SMCV or SMIC is selected as the optimal model as well. As a result, Tab. 4 shows the detection probability of each edge.

**Table 4:**
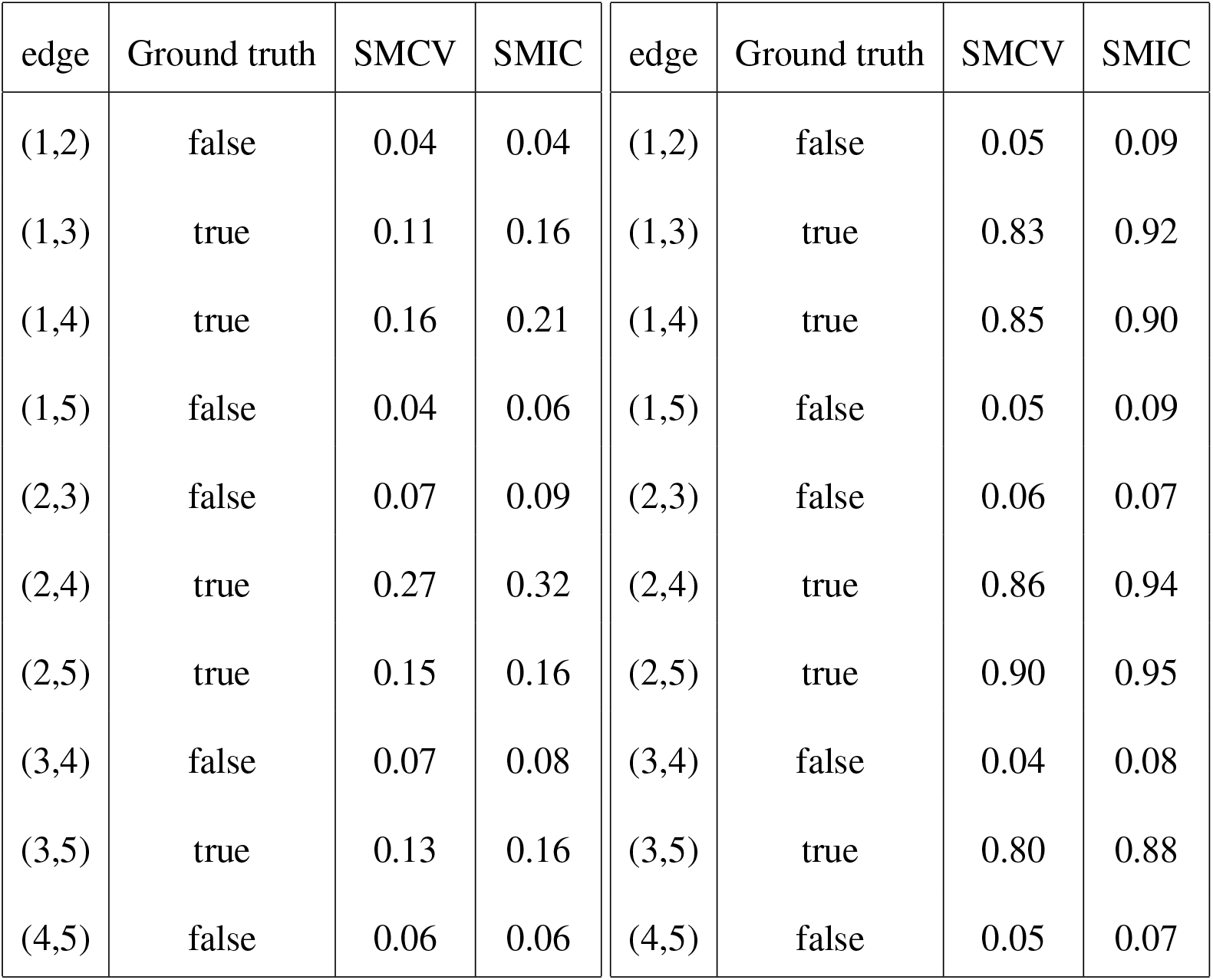
Detection Probability. The 5-dimensional synthetic data were sampled from the torus graph model with *c* = 0.1, 0.3.

### 3.3 Synthetic data generated from the Kuramoto model

Consecutively, we confirm the performance of our proposed method on synthetic data generated from the Kuramoto model (Kuramoto, 1975), which is a famous model to describe synchronization observed in the behavior of a large set of coupled oscillators such as neurons. To generate realistic synthetic data, we use the Kuramoto model with additional stochastic perturbations, involving five oscillators indexed by {1, 2, 3, 4, 5}:

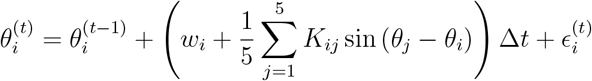

where *t* denotes the time step, Δ*t* denotes the time step size of discretization, *w*_*i*_ ∼ *N* (11.5, 2) is the intrinsic natural frequency of *i*-th oscillator that simulates the *α*-band, and the symmetric off-diagonal square matrix *K* denoting the undirected coupling effect. Here, we define the true graph structure as *G* = *G*(*V, E*), where *V* = {1, 2, 3, 4, 5} and *E* = {(1, 3), (1, 4), (2, 4), (2, 5), (3, 5)}, as in the previous experiment. The corresponding matrix *K* is defined as follows:

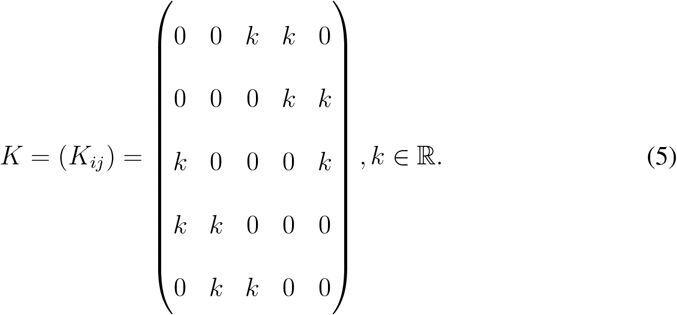

In addition to the original Kuramoto model, we introduce an additional noise term *ϵ*^(*t*)^, which is independently and identically distributed according to the von Mises distribution vM(0, 300), to make the generated data more realistic.

Figure 7 illustrates the successful performance of the proposed method in recovering the true graph structure, for example, in the case of *k* = 2. The number of synthetic data points was set to 25000, with Δ*t* = 0.01. In contrast, the traditional approach — computing PLV pairwise between nodes, followed by thresholding — struggles to accurately extract the true structure in this example. In contrast, our proposed method successfully minimizes the SMIC at the true graph structure, recognizing it as the best-fitting model and thereby enabling automatic model selection.

**Figure 7:**
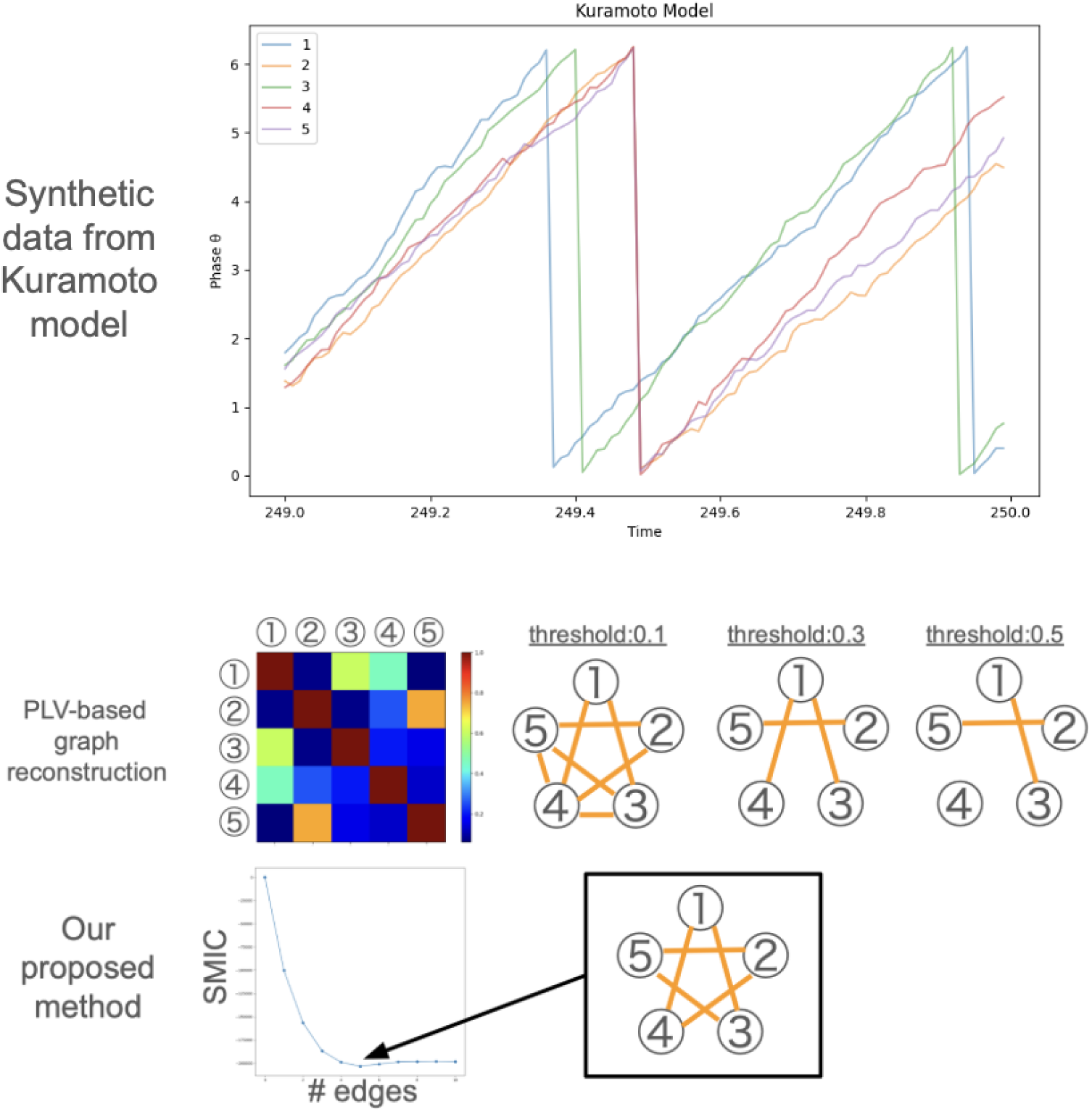
Proposed method performed on synthetic data generated from the Kuramoto model with 5 oscillators. *k* in Eq (5) is set to 2.

## 4 Real data application

In this section, we present two real-data applications, primarily to illustrate the utility of the proposed method for EEG/ECoG phase-based connectivity analysis.

### 4.1 Results on human EEG

By using a 61-dimensional human EEG dataset (50 seconds) collected from 20 participants, with seven labeled as the “drowsy” group and the rest as the “responsive” group based on their perceptual hit rate after administration of propofol, we investigated how differences in wakefulness modulate functional connectivity across different frequency bands. To investigate the dependence structures for each individual, we applied the rotational model with regularized score matching to each frequency band. This choice is supported by the experimental results that even when the full torus graph is applied to the real EEG data, the latter half of *ϕ*_*jk*_ in Eq (1) are always near zero, implying that the rotational model without these terms is sufficient for modeling these data. Fig. 8 shows one example. This is also similar to the model selection performed in the analyses using local field potentials as described by Klein et al. (2020).

**Figure 8:**
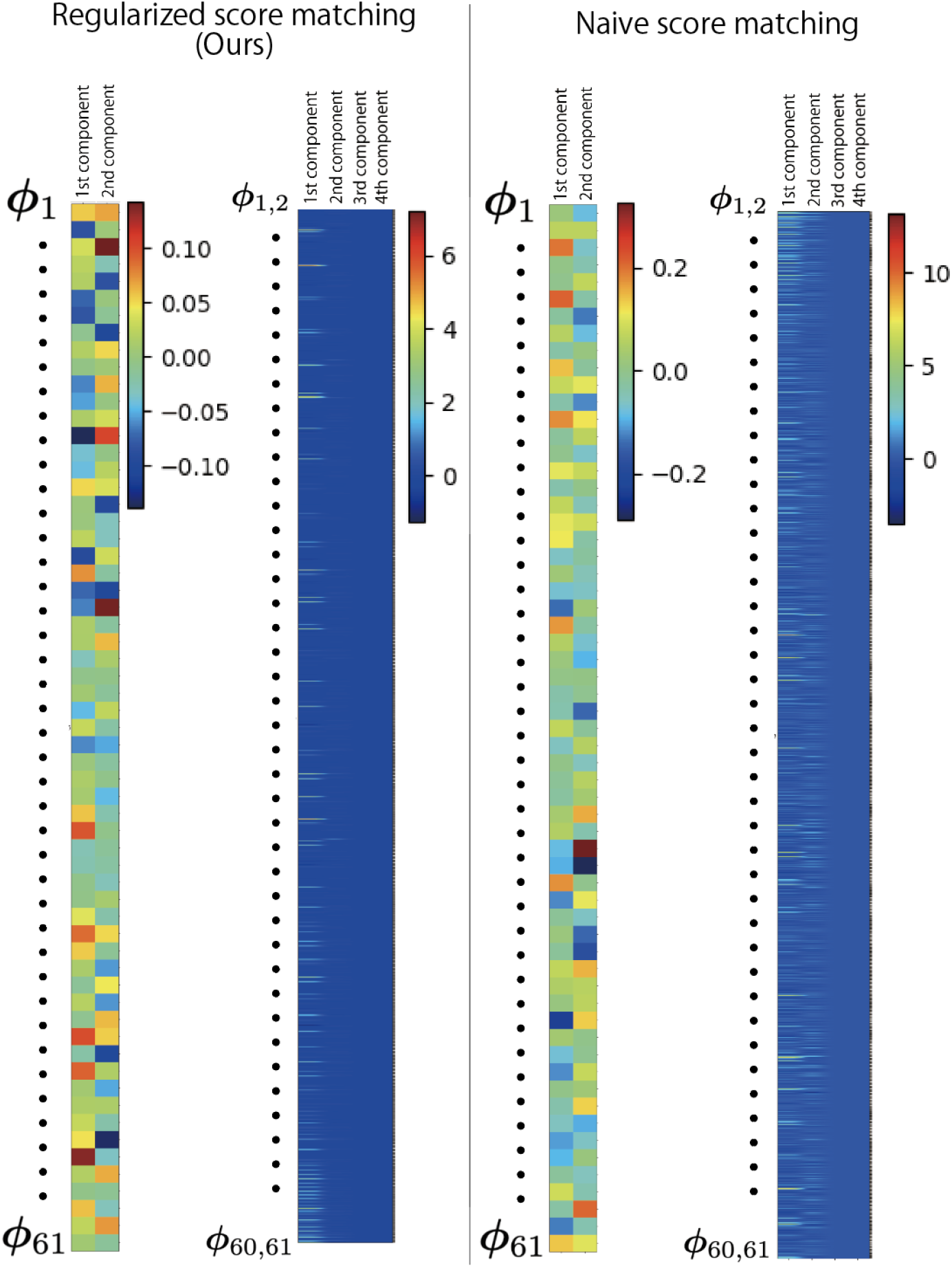
An example of the estimated values of 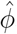. The left half shows the regularized score matching with Group LASSO and the right half shows the naive score matching. In each figure, the first image depicts *ϕ*_*i*_, 61 two-dimensional vectors, and the second image depicts *ϕ*_*jk*_, 1830 four-dimensional vectors, column by column.

The preprocessing is composed of the Hilbert transform and the bandpass filter extracting *α*-band (8Hz – 15Hz), *β*-band (12Hz – 25Hz), or *γ*-band (25Hz – 40Hz). These thresholds of frequency follow Chennu et al. (2016), who collected the dataset originally. In practice, the Hilbert transform is implemented by Python Scipy library. After the preprocessing (Hilbert transform and band pass filtering), we merge 10 epochs (0∼9) together to prepare 25000 samples. Then, we extract 61 dimensional data from it following the 10-10 system of electrodes (Koessler et al., 2009). The final data results in the shape of (*N, d*) = (25000, 61).

Fig. 9 displays one representative example from the drowsy group. The results are shown for the four participant state (baseline → mild → moderate → recovery), where the sparse solution is obtained for each state. For this participant, the number of edges varies at approximately 400 ∼ 600 edges, which is significantly smaller than the maximum edge number 61 · 60*/*2 = 1830. It is observed that the number of edges drops by the propofol injection between the baseline state and the mild state, while it recovers the network structure towards the recovery state gradually. To further enhance the interpretability of these networks, observing the underlying community structure of them is an useful approach. Here, the color of each node represents the optimal community structure obtained by the modularity maximization (Newman and Girvan, 2004), where the number of communities is not specified in advance but determined through the optimization process. Overall, it is observed that the close nodes form a cluster in general, while some clusters are dispersed among the whole brain, e.g., the gray-colored cluster in the mild state. These community structures depict the occurrence of neural oscillations with richer information compared to the naive way of calculating the correlation between each node pairs.

**Figure 9:**
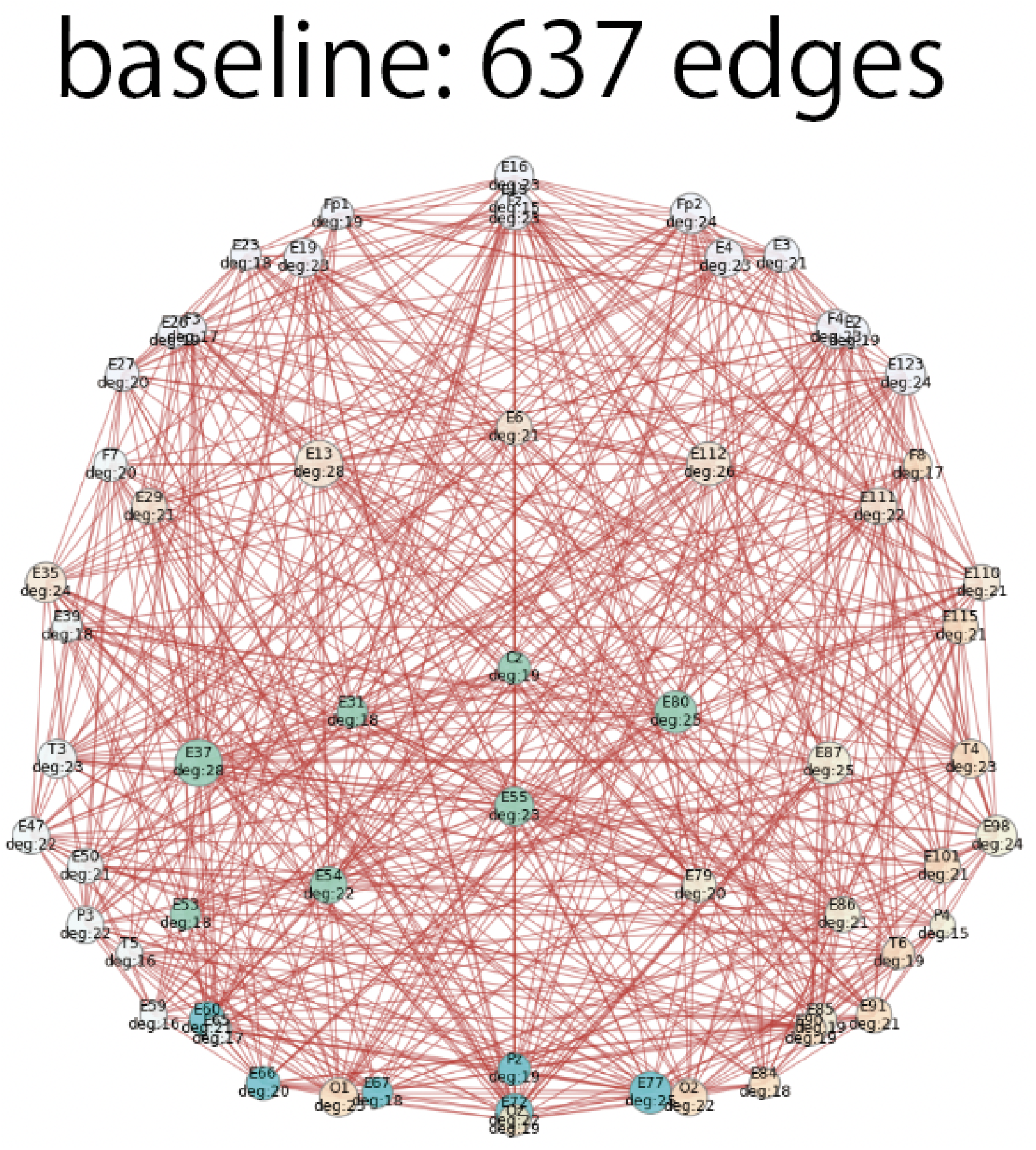
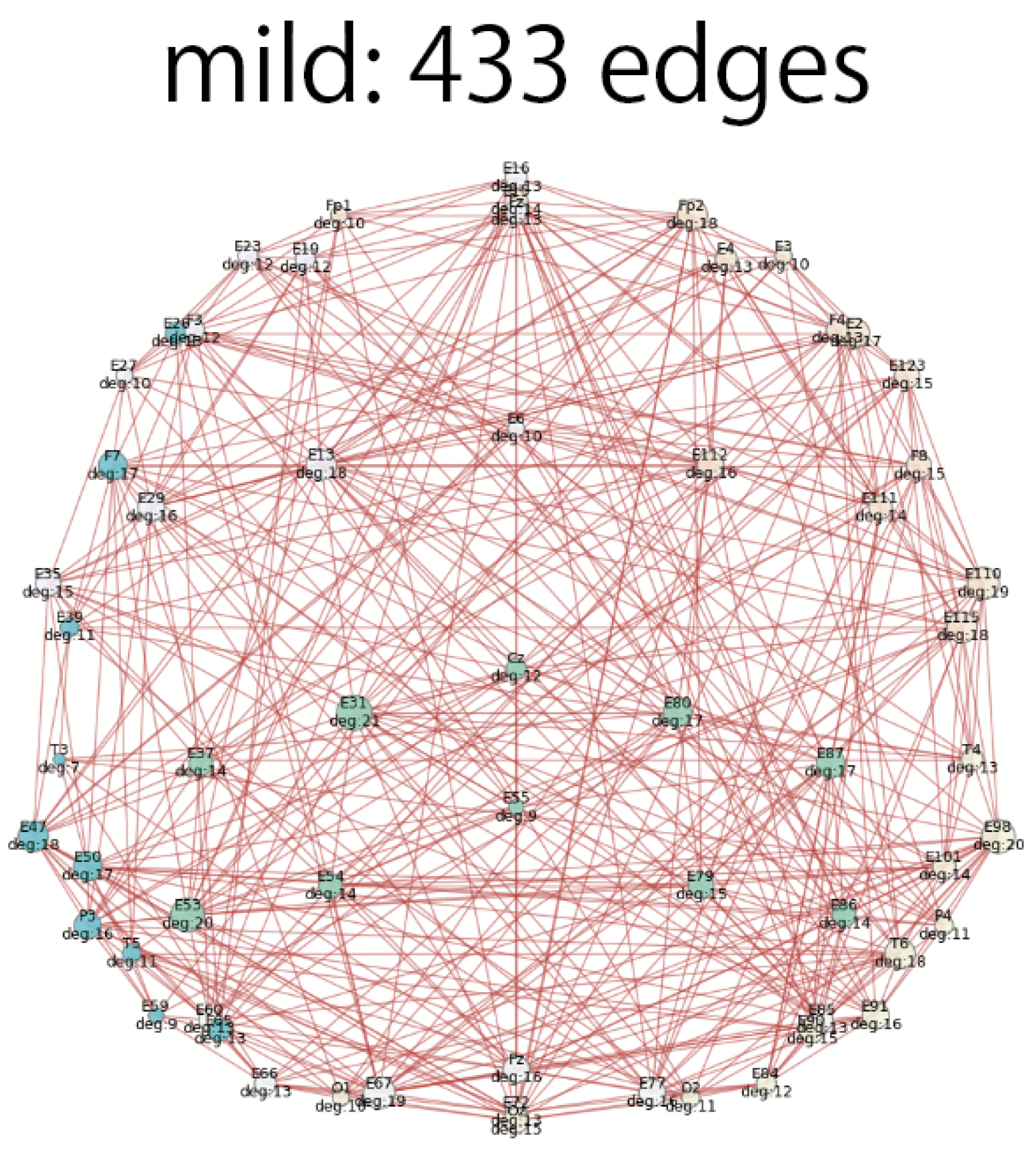
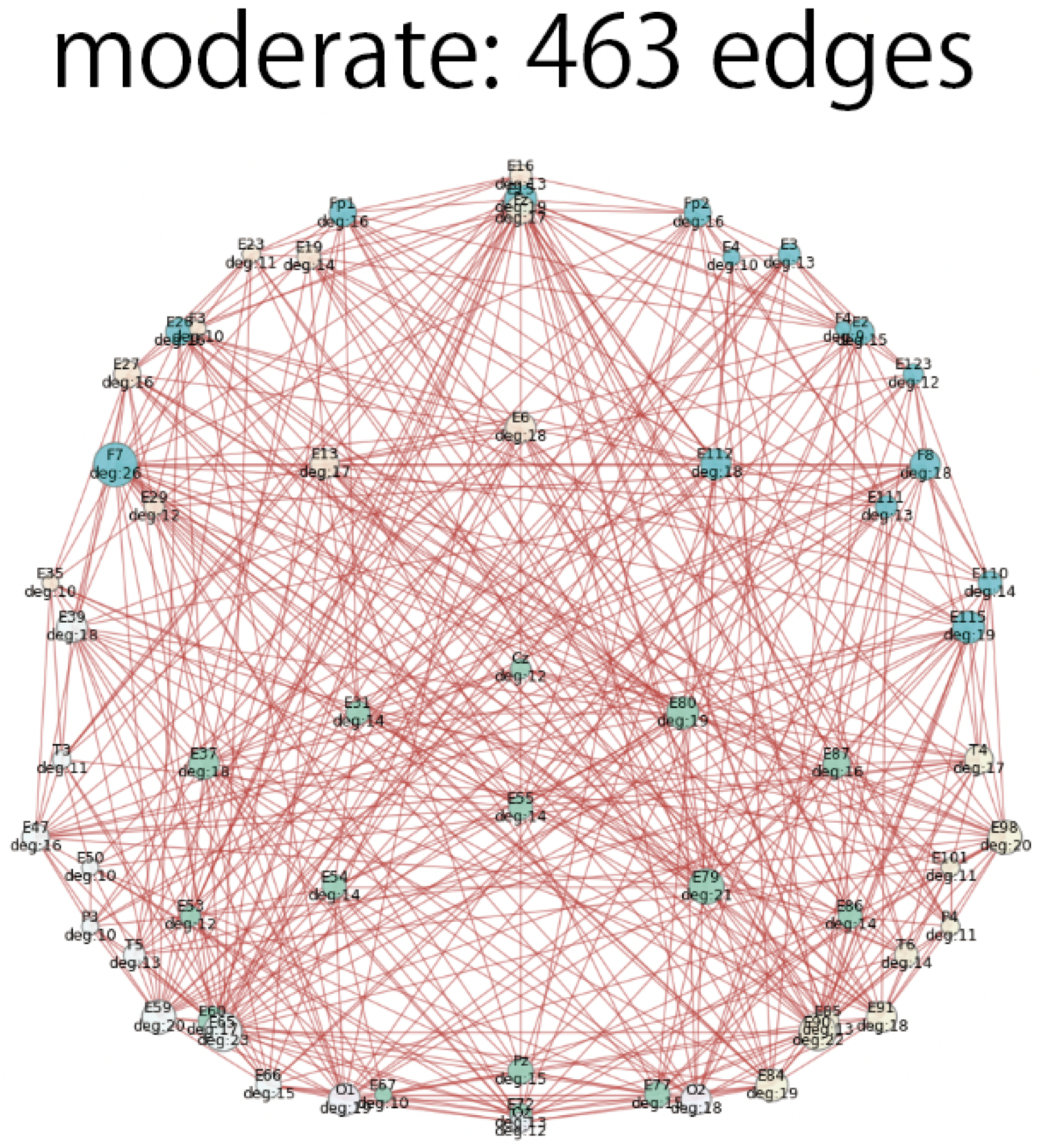
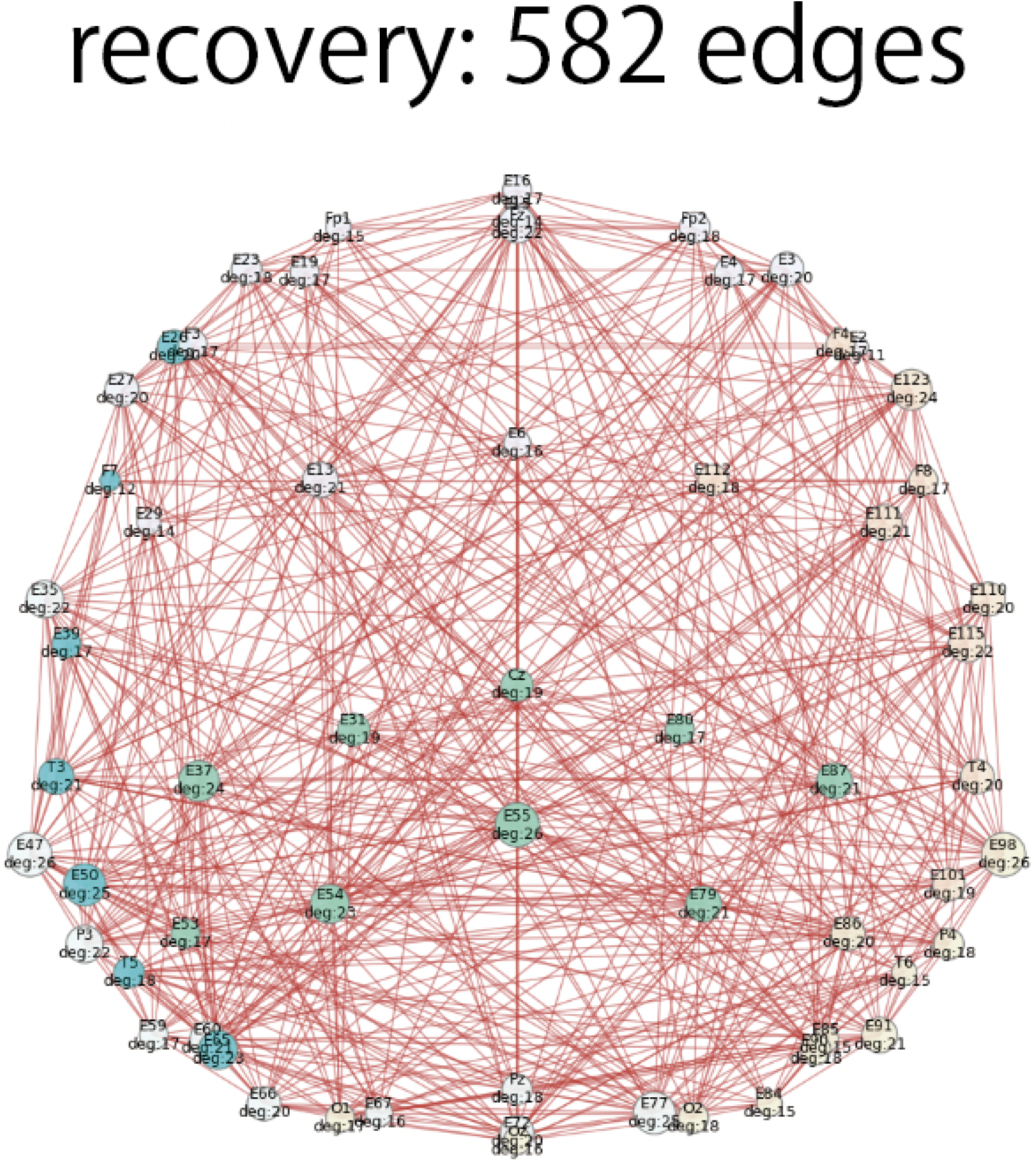
Change in reconstructed graph structures. (baseline → mild → moderate → recovery; participant_ID = 7 from the drowsy group). Node colors represent community structures identified through modularity maximization.

For all participants, Tab. 5 and Tab. 6 summarize the changes in the number of edges, modularity, average clustering coefficient, and average shortest path length for the recovered graph structures for each band of EEGs. The shifts of the ratio between average clustering and average shortest path length during each state are shown in Fig. 10. We also present uncertainty analysis by permutation tests associated with these results in Tab. 7 and Tab. 8. Here, to assess whether the group means differ, we randomly permuted the group labels 10000 times and recomputed the mean difference each time. The p-value was calculated as the proportion of permuted mean differences whose absolute value was at least as large as the observed mean difference. As a result, the responsive group clearly exhibited higher values in every frequency band analyzed. A significant distinction between the drowsy group and the responsive group cannot be observed in the alpha band EEGs. Contrarily, for the beta bands, the drowsy group tends to exhibit a lower number of edges, higher modularity, lower average clustering coefficient, and higher average shortest path length compared to the responsive group. Given that a small-world network (Watts and Strogatz, 1998; Muldoon et al., 2016) is primarily characterized by a higher average clustering coefficient and shorter average shortest path (Humphries and Gurney, 2008), we conclude that small-worldness is more pronounced in the beta connectivity of the responsive group, while modularity is not increased in this group. This is not aligned with the previous study on 10 healthy humans (Lee et al., 2013), which reported that propofol increased both average path length, clustering coefficient, and modularity calculated from phase lag index (PLI) network. This discrepancy could be due to several factors, including differences in the methodologies used, the specific experimental conditions, or the statistical method for reconstructing the graph network structure from the EEG data.

**Table 5:** Summary of graphical properties (the number of edges and modularity). The average is taken among all the participants in each participant group. Refer to S4 Appendix for full individual results.

|  | participant group | baseline | mild | moderate | recovery |
| --- | --- | --- | --- | --- | --- |
| $\alpha$ band (8-15Hz) | drowsy | (762, 0.388) | (855, 0.353) | (749, 0.394) | (863, 0.351) |
| $\alpha$ band (8-15Hz) | responsive | (863, 0.364) | (865, 0.346) | (840, 0.358) | (886, 0.348) |
| $\beta$ band (12-25Hz) | drowsy | (844, 0.411) | (857, 0.393) | (788, 0.421) | (944, 0.376) |
| $\beta$ band (12-25Hz) | responsive | (870, 0.399) | (980, 0.361) | (967, 0.358) | (953, 0.368) |
| $\gamma$ band (25-40Hz) | drowsy | (836, 0.424) | (795, 0.423) | (799, 0.428) | (772, 0.444) |
| $\gamma$ band (25-40Hz) | responsive | (851, 0.402) | (995, 0.359) | (985, 0.353) | (962, 0.366) |

**Table 6:** Summary of graphical properties (average clustering coefficient and average shortest path length). The average is taken among all the participants in each participant group. Refer to S4 Appendix. for full individual results.

|  | participant group | baseline | mild | moderate | recovery |
| --- | --- | --- | --- | --- | --- |
| $\alpha$ band (8-15Hz) | drowsy | (0.427, 1.583) | (0.272, 0.881) | (0.156, 0.574) | (0.125, 0.403) |
| $\alpha$ band (8-15Hz) | responsive | (0.478, 1.529) | (0.259, 0.823) | (0.162, 0.535) | (0.126, 0.389) |
| $\beta$ band (12-25Hz) | drowsy | (0.475, 1.539) | (0.275, 0.876) | (0.166, 0.565) | (0.137, 0.391) |
| $\beta$ band (12-25Hz) | responsive | (0.487, 1.524) | (0.289, 0.791) | (0.185, 0.511) | (0.136, 0.380) |
| $\gamma$ band (25-40Hz) | drowsy | (0.467, 1.544) | (0.262, 0.894) | (0.166, 0.564) | (0.120, 0.421) |
| $\gamma$ band (25-40Hz) | responsive | (0.478, 1.535) | (0.293, 0.787) | (0.189, 0.507) | (0.138, 0.378) |

**Table 7:** Permutation test p-values for network characteristics (edges, modularity, clustering coefficient, shortest path length) between two different patient conditions. The three tables correspond to *α*-, *β*-, and *γ*-band from top to bottom. Bold values denote *p <* 0.05.

|  | baseline | mild | moderate | recovery |
| --- | --- | --- | --- | --- |
| baseline | — | (0.448, 0.070, 0.296, 0.525) | (0.552, 0.891, 0.679, 0.509) | (0.105, **0.011**, 0.077, 0.105) |
| mild | — | — | (0.148, 0.052, 0.113, 0.177) | (0.618, 0.952, 0.707, 0.537) |
| moderate | — | — | — | **(0.029, 0.037, 0.034, 0.026)** |
| recovery | — | — | — | — |

|  | baseline | mild | moderate | recovery |
| --- | --- | --- | --- | --- |
| baseline | — | **(0.059, 0.011, 0.069, 0.060)** | (0.263, **0.046**, 0.219, 0.278) | **(0.022, 0.001, 0.020, 0.022)** |
| mild | — | — | (0.300, 0.395, 0.402, 0.289) | (0.806, 0.928, 0.776, 0.799) |
| moderate | — | — | — | (0.327, 0.427, 0.319, 0.320) |
| recovery | — | — | — | — |

|  | baseline | mild | moderate | recovery |
| --- | --- | --- | --- | --- |
| baseline | — | (0.069, **0.019, 0.042**, 0.075) | **(0.029, 0.005, 0.017, 0.030)** | (0.288, 0.198, 0.115, 0.524) |
| mild | — | — | (0.871, 0.782, 0.857, 0.892) | (0.498, 0.277, 0.697, 0.366) |
| moderate | — | — | — | (0.551, 0.157, 0.753, 0.420) |
| recovery | — | — | — | — |

**Table 8:** Permutation test p-values for network characteristics (edges, modularity, clustering coefficient, shortest path length) between two different patient groups. The three tables correspond to *α*-, *β*-, and *γ*-band from top to bottom. Bold values denote *p <* 0.05.

| Phase | drowsy vs responsive |
| --- | --- |
| baseline | (0.090, 0.300, 0.100, 0.090) |
| mild | (0.911, 0.842, 0.987, 0.882) |
| moderate | (0.184, 0.283, 0.264, 0.172) |
| recovery | (0.717, 0.908, 0.636, 0.715) |
| Phase | drowsy vs responsive |
| baseline | (0.717, 0.667, 0.747, 0.710) |
| mild | (0.101, 0.308, 0.125, **0.099**) |
| moderate | (**0.017**, **0.033**, **0.036**, **0.016**) |
| recovery | (0.898, 0.736, 0.857, 0.900) |
| Phase | drowsy vs responsive |
| baseline | (0.814, 0.419, 0.730, 0.809) |
| mild | (**0.006**, **0.035**, **0.010**, **0.006**) |
| moderate | (**0.001**, **0.006**, **0.003**, **0.001**) |
| recovery | (**0.044**, **0.006**, 0.057, **0.034**) |

**Figure 10:**
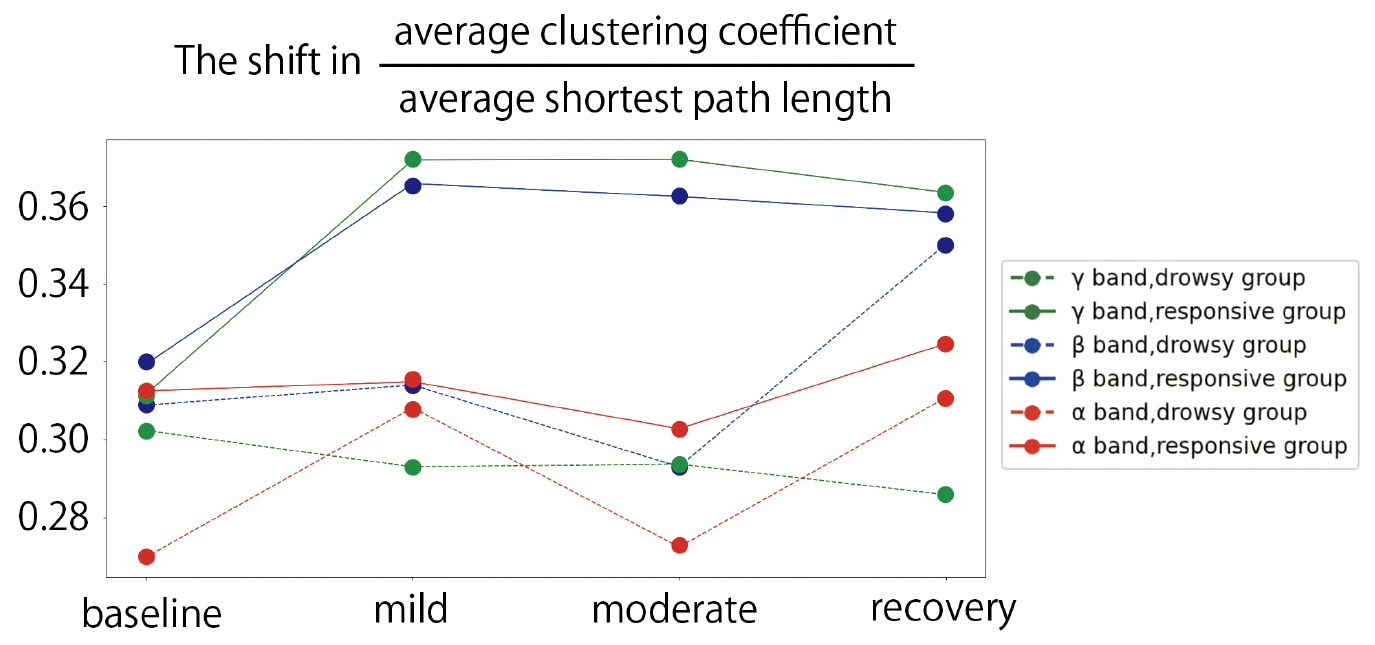
The shift in graphical properties during each participant state. The ratio between the average clustering coefficient and the average shortest path length in each participant state.

Additionally, we observe the relationship between different frequency bands since the cross-frequency coupling has been crucial in EEG analysis. Fig. 11 shows pairwise scatter plots of modularity across frequency bands. For each EEG recording, we band-pass filter the signal into the *α, β*, and *γ* bands, reconstruct a connectivity graph for each band using our method, and compute its modularity. Each point therefore corresponds to a single recording, with coordinates given by the modularity of the two bands being compared (e.g., *β* on *x*-axis vs. *γ* on *y*-axis). The responsive group exhibits a significantly stronger *β*-*γ* correlation than the drowsy group. This difference in patterns may potentially serve as a feature for predicting the participant group solely from EEG recordings, without the need to conduct the injection experiment.

**Figure 11:**
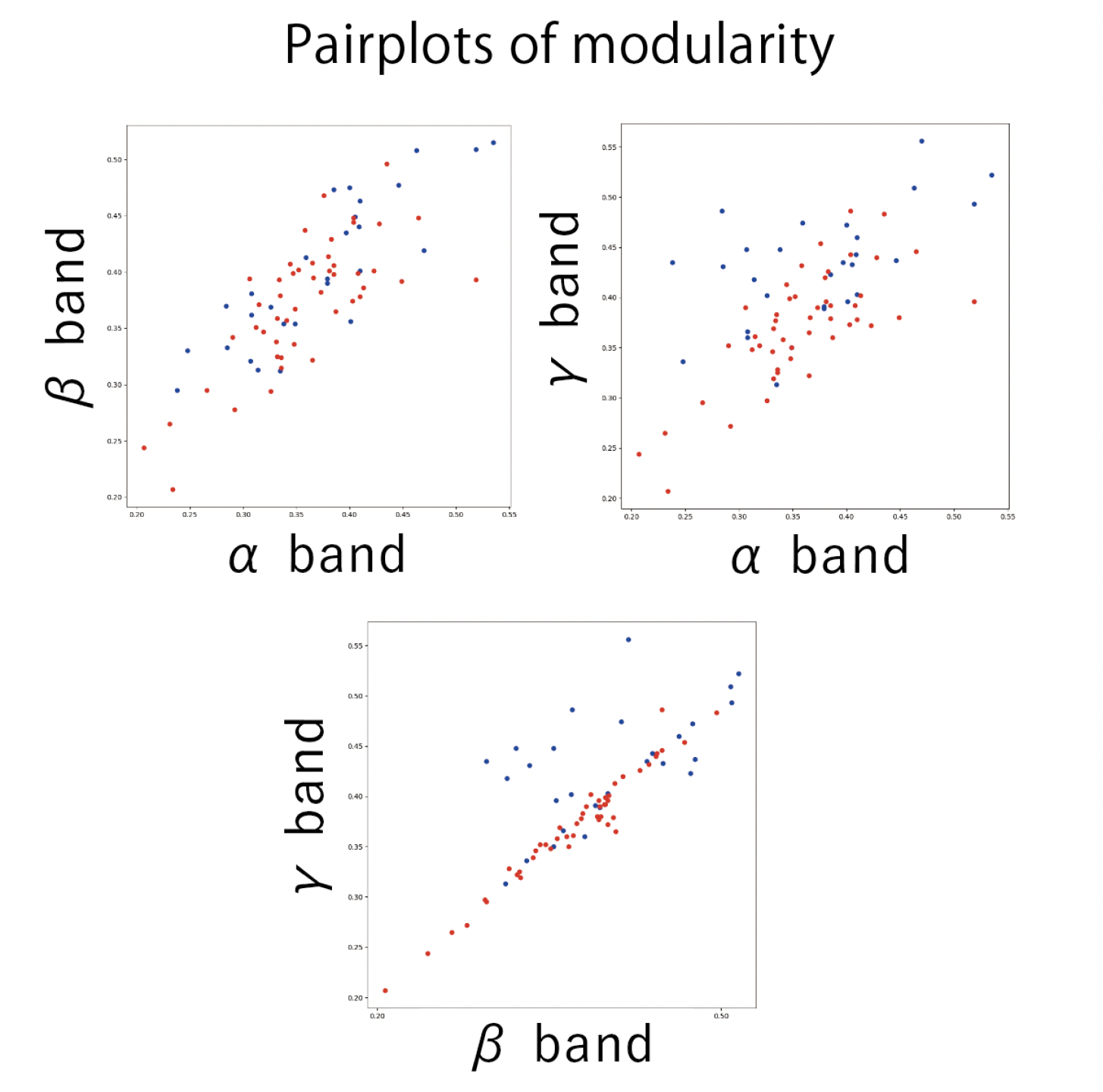
Pairplots of modularity from different frequency bands. From each EEG recording, *α, β*, and *γ* bands were extracted by bandpass filtering; a graph was recovered for each band and its modularity computed. Scatter plots are shown for *α*–*β, α*–*γ*, and *β*–*γ*; each dot represents one recording, positioned by the modularity values of the two bands. The blue points are from the drowsy group whereas the red points are from the responsive group. The responsive group shows a markedly stronger *β*–*γ* correlation than the drowsy group.

### 4.2 Results on marmoset ECoG

To further apply the proposed method for visualizing the phase-based connectivity, we used the 96 dimensional ECoG series data of marmosets. According to the original paper (Komatsu and Ichinohe, 2021), these data were collected from two individuals — one recorded from the left cerebral hemisphere and the other from the right. For example, the *α* bands of resting-state ECoG from marmosets were extracted and depicted in Fig. 12. For the data taken from the left hemisphere, the SMIC is minimized at the model with 680 edges. Contrarily, for the data taken from the right hemisphere, the SMIC is minimized at the model with 618 edges. In both cases, considering that the maximum number of edges can be as high as _96_C_2_ = 4560, the edges in the resulting graph are kept relatively sparse. The color of nodes represents the optimal community structure obtained by modularity maximization. Thus, due to the regularization effect of the proposed method, a sparse network structure with clearly defined community structure is obtained as a result.

**Figure 12:**
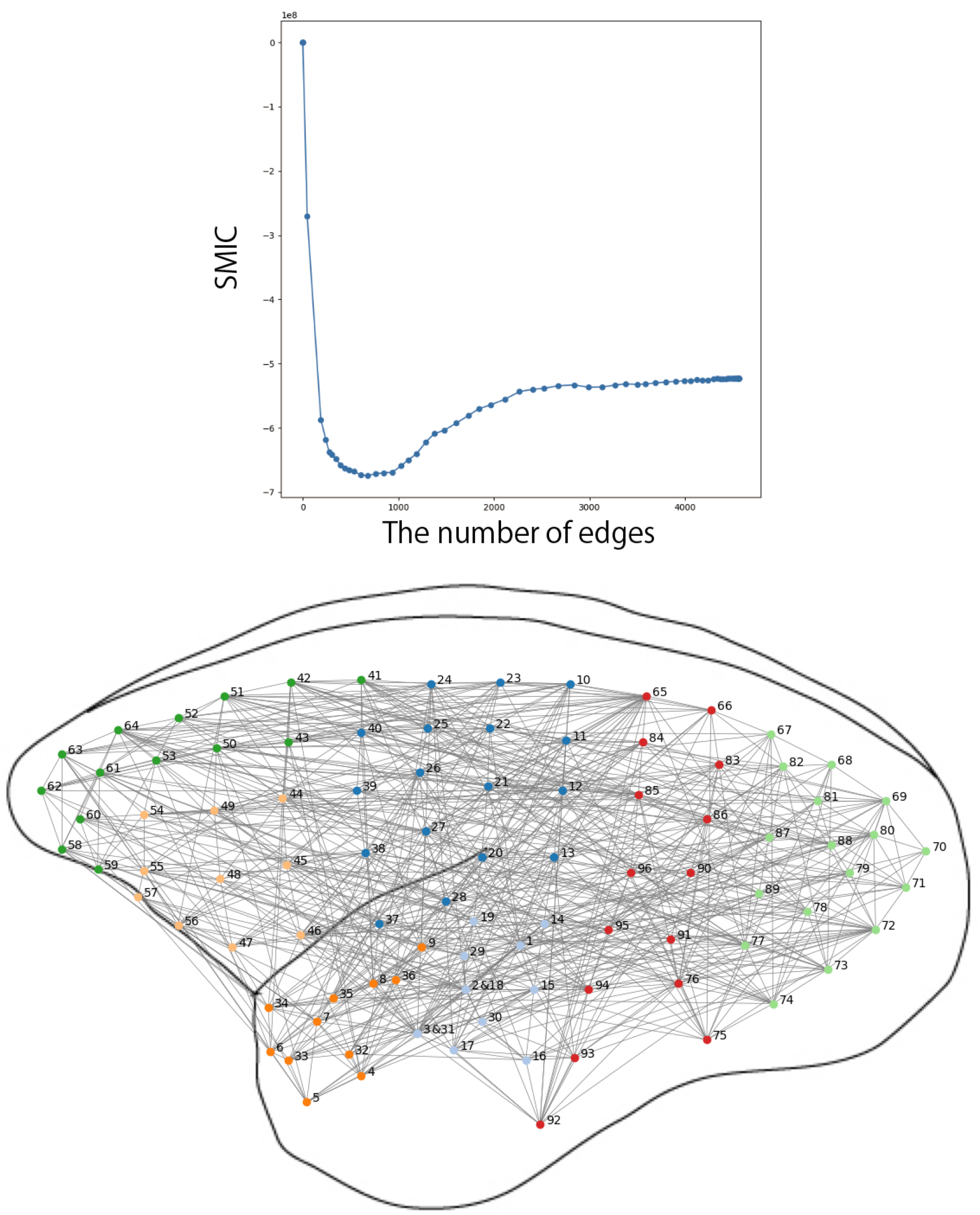
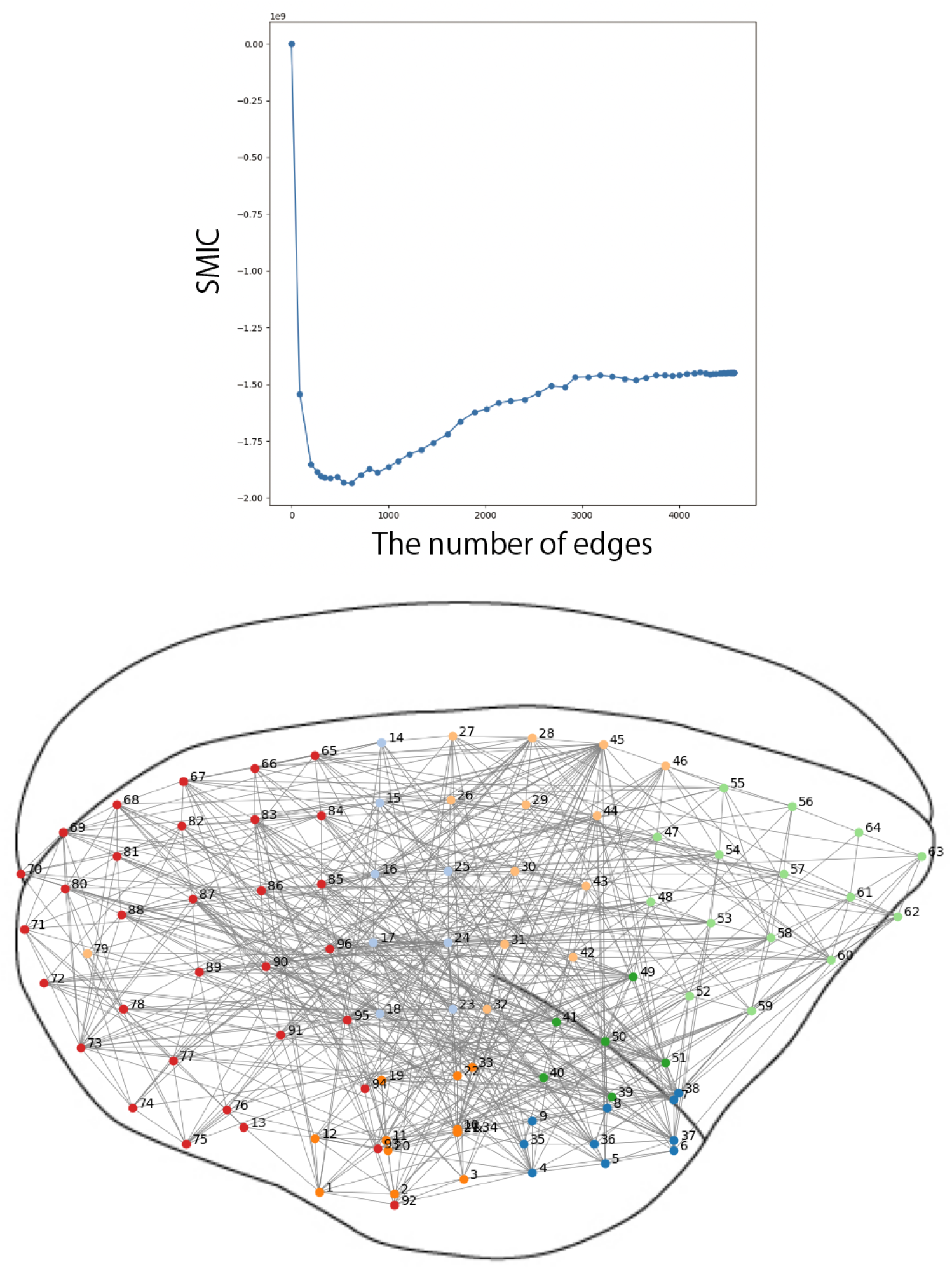
Results of the proposed method applied to the *α* band of two resting-state marmoset ECoG datasets. Axes in blue graphs are labeled with SMIC (y-axis) and the number of edges (x-axis).

To observe the shift in the data between before and after the ketamine injection, the number of edges and modularity in each state are summarized in Tab. 9. The network structure becomes less modular for the first marmoset, whereas the opposite is true for the second marmoset. These observations, based on the comparison of network modularity, are interesting in the context of previous studies that have explored the effects of anesthetics, including ketamine, on brain network architecture.

**Table 9:** The shift in the recovered network structure from marmoset ECoG.

| Marmoset ID | R03_0024_CM622M | R03_0035_CM814F |
| --- | --- | --- |
| Target brain | left hemisphere | right hemisphere |
| Resting-state ($\alpha$ band) | #edge = 680, modularity = 0.634 | #edge = 618, modularity = 0.594 |
| after 10 min. ($\alpha$ band) | #edge = 695, modularity = 0.600 | #edge = 505, modularity = 0.622 |
| Resting-state ($\beta$ band) | #edge = 435, modularity = 0.666 | #edge = 379, modularity = 0.623 |
| after 10 min. ($\beta$ band) | #edge = 798, modularity = 0.595 | #edge = 368, modularity = 0.648 |
| Resting-state ($\gamma$ band) | #edge = 370, modularity = 0.673 | #edge = 360, modularity = 0.655 |
| after 10 min. ($\gamma$ band) | #edge = 576, modularity = 0.631 | #edge = 321, modularity = 0.657 |

Many prior studies based on EEG signals in humans have consistently shown that from the awake to anesthetized state, the brain’s functional connectivity is reduced (Hudetz, 2012). In contrast, some ECoG studies in monkeys report that the overall functional connectivity shows an increasing trend in anesthetized state (Xie et al., 2021). For instance, using phase-amplitude coupling to build functional connectivity, propofol-induced anesthesia enhances frontal and parietal interactions in monkeys (Ma et al., 2019). These contrasting findings could reflect the complex and dynamic nature of ketamine’s interaction with neural networks, where individual differences in brain structure or the specific nature of the intervention could yield divergent outcomes in terms of network topology. However, compared to modularity analysis based on EEG sig-nals (Kim et al., 2018), modularity analysis based on ECoG signals is still limited. This is an important area for future work, as the network topology outcomes could depend on the target animal and the type of signal used.

The changes in network modularity we observed were consistent across all frequency bands, adding an important layer of insight. Previous studies (e.g., Yan et al. (2022)) have demonstrated that different frequency bands of ECoG in monkeys respond in opposite ways to acute ketamine injection. Our results, showing that the modularity changes occurred across all frequency bands, suggest that ketamine has a global effect on brain network topology, influencing both low and high-frequency oscillations. This finding contrasts with previous research that has highlighted frequency-specific effects of ketamine, indicating that the drug’s impact on network dynamics may involve more complex, widespread alterations in brain activity.

Although the number of marmoset samples is too low to draw any solid conclusions, our results suggest the possibility of predicting an individual’s state based on network modularity extracted from ECoG. Considering the increasing use of ketamine in clinical contexts, understanding the individual variability in ketamine’s effects on brain network topology could have significant implications for personalized treatment strategies. It also emphasizes the need for further studies with larger sample sizes to better understand ketamine’s impact on brain networks.

## 5 Discussion and Conclusion

Through two synthetic datasets, as well as real EEG and ECoG examples, we have demonstrated the effectiveness of our proposed method, which fits a torus graph to the observed data with regularization to yield a sparse network structure. Although regularization in high-dimensional sparse graphical modeling is not new — for example, in pairwise interaction models (Lin et al., 2016) — its application to circular data, such as EEG/ECoG phase, represents a novel contribution. Especially when compared with conventional methods that threshold correlation or PLV for node-wise pairs individually, our method estimates the entire network structure simultaneously. Most importantly, it is free from arbitrary binarization and is supported by a solid statistical foundation. It is also free from cross validation, which is computationally expensive and typically required when solving LASSO-like problems. These are made possible because SMIC minimization (Matsuda et al., 2021) directly determines the graph struc-ture that best fits the data.

We acknowledge a few limitations in our proposed method. Since our proposed method is based on torus graph modeling, it inevitably inherits the inherent limitations of this model. First, higher-order interactions or cross-frequency couplings are not captured, as the torus graph model is limited to linear combinations of sine and cosine terms as in Eq (1). This restricts the model’s ability to represent more complex or nonlinear phase interactions that may be present in real neural systems. Second, our method disregards temporal dependencies in the phase data. That is, the analysis is conducted on a static snapshot of the data, without modeling how interactions evolve over time. While this simplification facilitates computational efficiency and enables principled statistical estimation, it may reduce the method’s applicability to dynamically changing systems, which are likely to be prevalent in the brain. In contrast, prior work in the physics literature has focused on reconstructing temporal phase dynamics to infer directional coupling between oscillators based on their time series (Rosenblum and Pikovsky, 2001; Kralemann et al., 2011). These methods aim to reconstruct directed, time-resolved interactions, typically under the assumption of well-defined oscillatory behavior. However, such approaches often rely on strong model assumptions or adhoc procedures and may lack a statistical framework for model selection or uncertainty quantification. Our method, which is non-directional in nature, offers a complementary perspective by enabling the estimation of global graph structure in a statistically grounded way, without requiring explicit time-series modeling or prior knowledge of oscillator dynamics. Addressing these limitations would require extending the model to incorporate nonlinear features, multi-frequency dynamics, or temporal structure, which remains an important direction for future work.

In summary, we extended the score matching estimation method for the torus graph model proposed by Klein et al. (2020) by applying Group LASSO-type regularization to achieve a sparse solution, enhancing the interpretation of EEG/ECoG phase-based connectivity analysis. Our method, implemented via the ADMM algorithm and SMIC minimization, was validated through simulation studies using synthetic data sampled from the prespecified torus graph model and the Kuramoto model. We also applied our method to real EEG/ECoG data analysis and found that it contributes to a more sophisticated approach for analyzing high-dimensional phase-based functional connectivity, despite certain limitations, such as the disregard for time series structure including autocorrelations. From the human EEG dataset, we found that the modularity of the reconstructed brain network structure exhibits different tendencies when the *β* and *γ* bands are used; the modularity of *β* band connectivity is nearly the same as that of *γ* band connectivity in the responsive group, whereas the latter tends to have a larger value than the former in the drowsy group. Similar analyses were applied to marmoset ECoG data with ketamin injection, as our method is domain-agnostic.

Future research could focus on spatio-temporal analysis by further incorporating time series features and the relative positions of each electrode. Additionally, exploring the integration of our method with the source localization problem may yield valuable insights into brain connectivity.

## Acknowledgments

Issey Sukeda was supported by RIKEN Junior Research Associate program. Takeru Matsuda was supported by JSPS KAKENHI Grant Numbers 22K17865, 24K02951, and JST Moonshot Grant Number JPMJMS2024. We also thank Dr. Yoshihito Saito and Dr. Sai Tanimoto for supportive comments.

## Appendix

### S1 Appendix. Computational time of naive score matching (top) / Group LASSO (bottom) in seconds

In Tab. 10, the latter includes the former once per run because calculating SMIC requires the naive score matching estimator. Since our method is based on LASSO algorithm, it is expected to work in reasonable computational time even for high-dimensional problems. Specifically, when the dimension of data is 61, the full torus graph model includes 7442 parameters and the rotational model includes 3782 parameters.

**Table 10:**
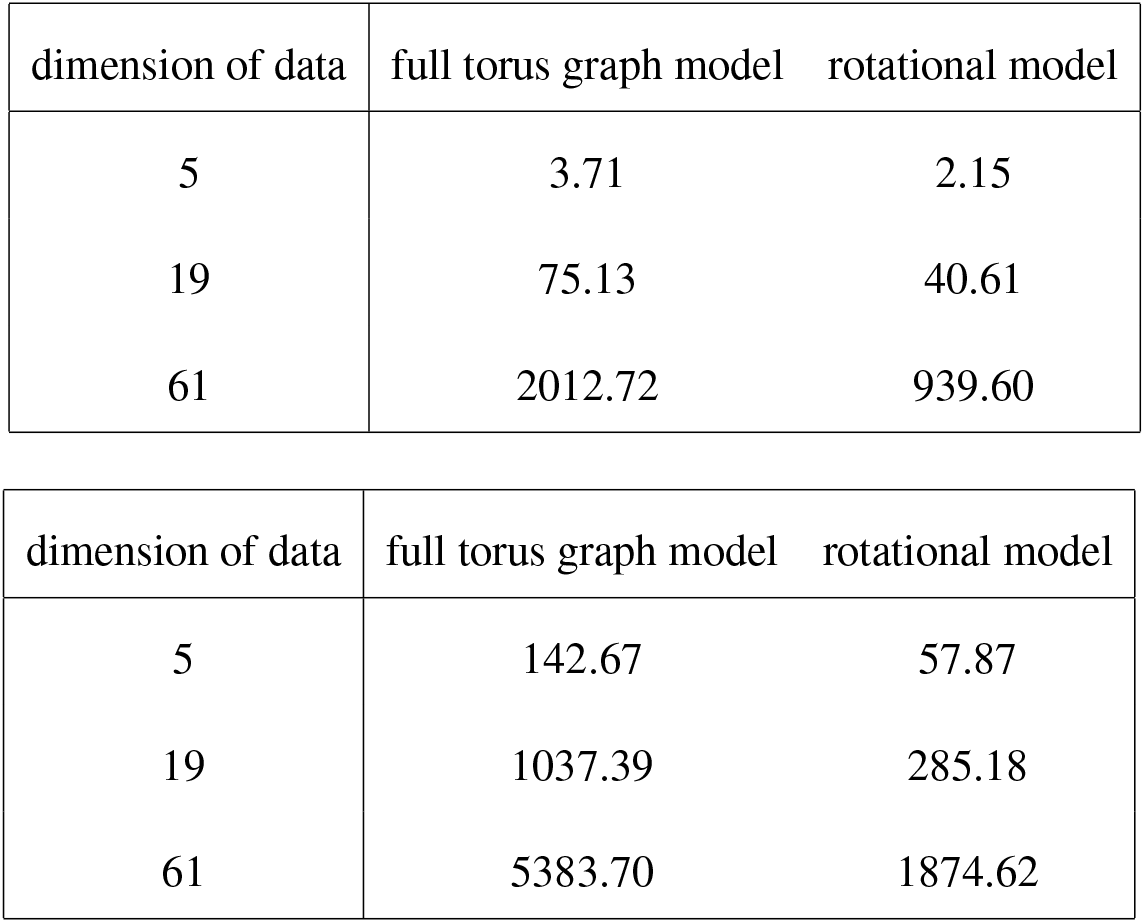
Computational time of naive score matching (top) / Group LASSO (bottom) in seconds.

### S2 Appendix. ADMM : the optimization algorithm for calculating score matching estimators

Since the score matching estimator with regularization term have the similar objective function to the LASSO problem, ADMM algorithm can be applied to calculate it. The following Algorithm 1 and 2 describe the whole process.

#### Algorithm 1 ADMM (for Group LASSO)

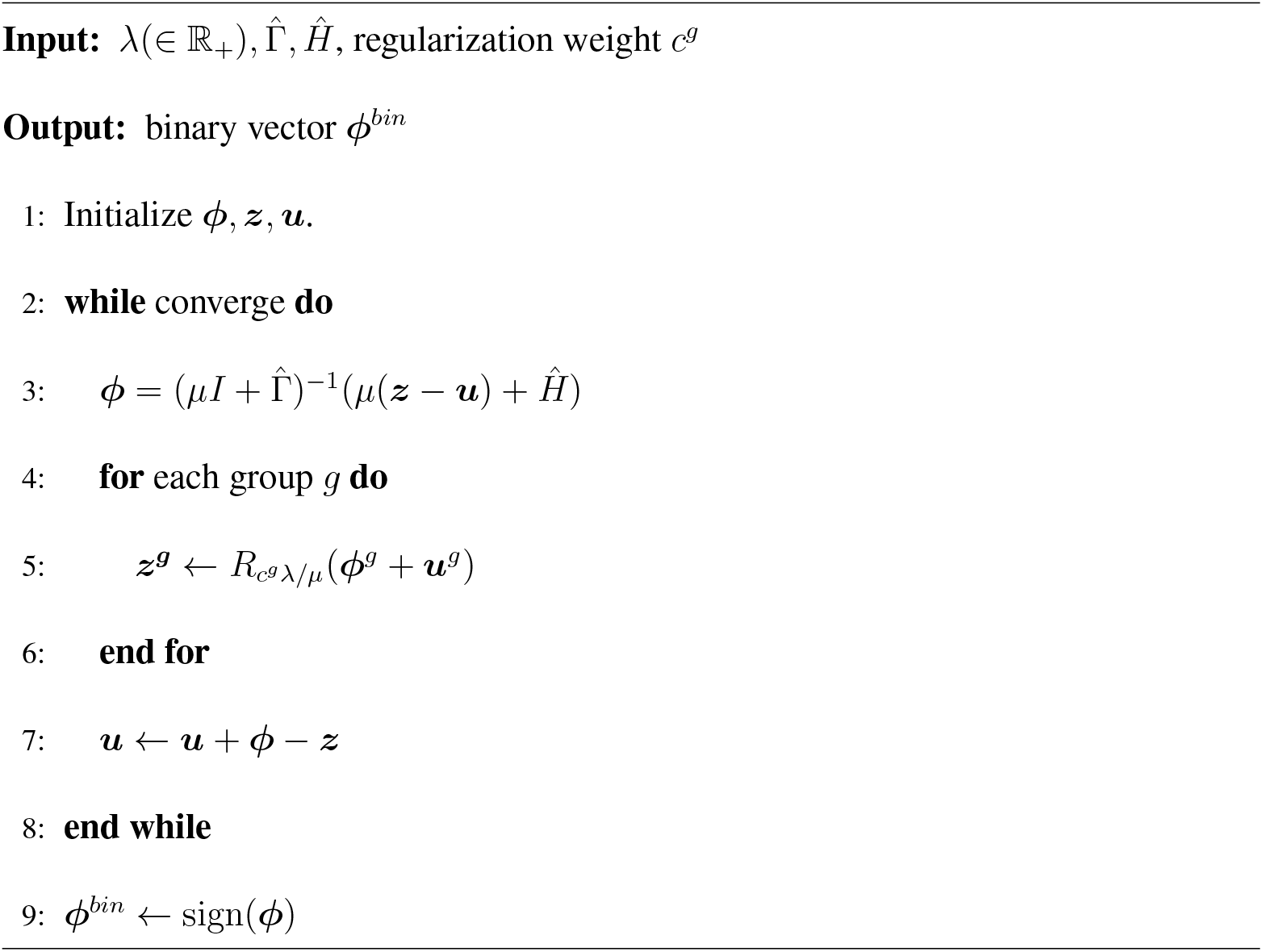

#### Algorithm 2 Regularized score matching of the torus graph via SMIC minimization

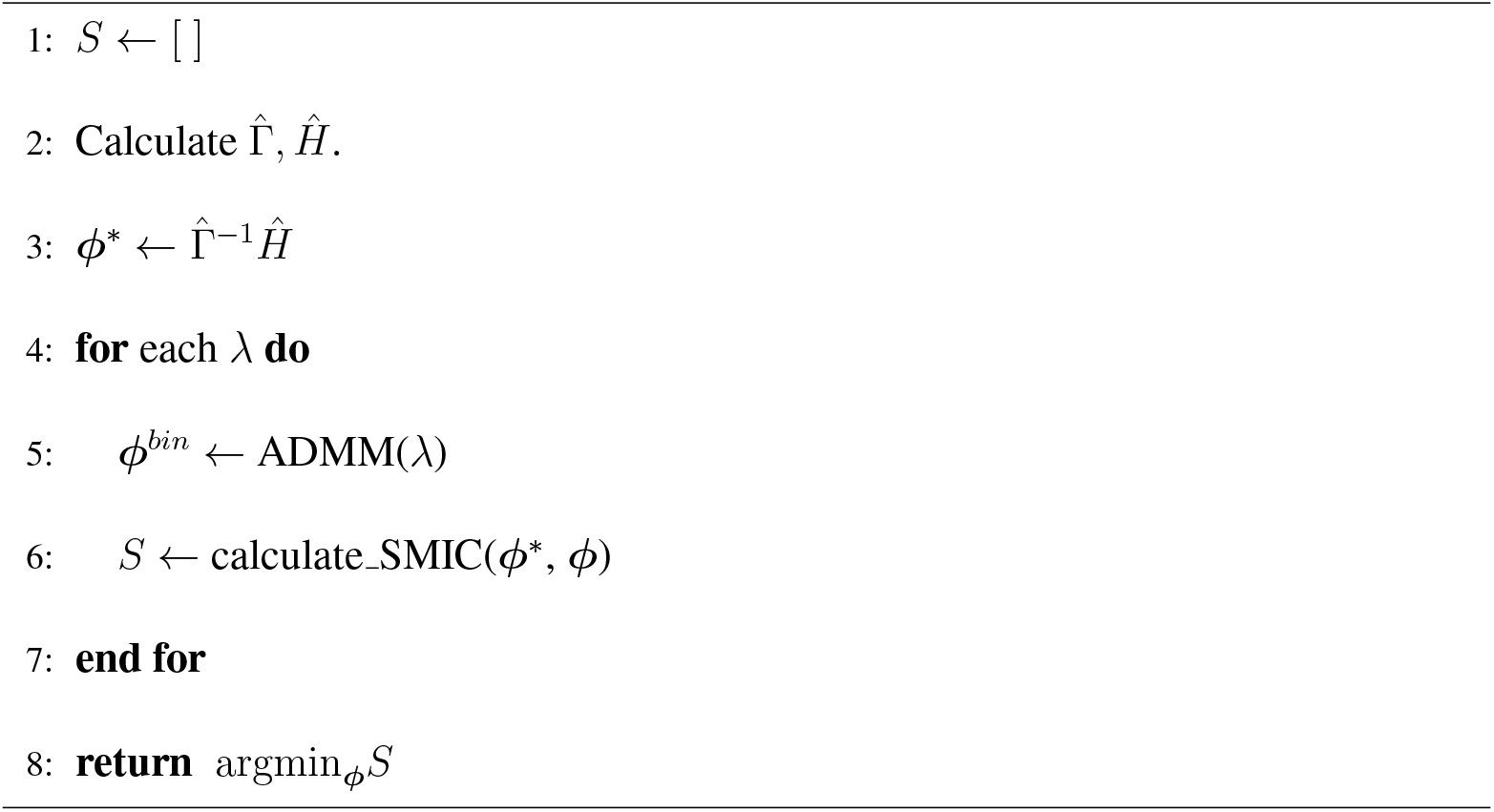

### S3 Appendix. Bias correction role of the SMIC

In general, information criteria is constructed by correcting the negative bias of the discrepancy between the empirical distribution using samples and the plug-in model (predictive distribution) when estimating the discrepancy between the true distribution and the model. The predictive distribution via score matching estimator is obtained as 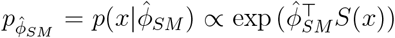.The score matching objective 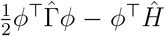 is an unbiased estimator of 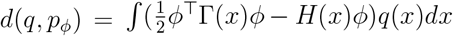, where *q*(*x*) is the unknown true distribution. However,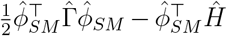 has an inherit bias when used for estimating 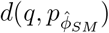. The true bias can be calculated as

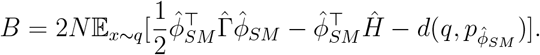

On the other hand, the bias correction term of SMIC is an unbiased estimator of this bias:

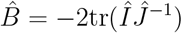

as in (4).

Here we confirm that 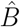 is truly an unbiased estimator of *B* by numerical simulations. We use the mixture of two bivariate von Mises distribution (Mardia, 1975) (2-dimensional torus graph) as the true distribution:

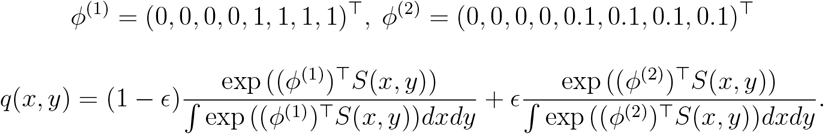

The first distribution is more concentrated to (0, 0), whereas the second distribution has larger variance and functions as adding a noise.

We set *N* = 1000. The values *B* and 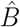 were computed by the Monte Carlo simulation with 10^5^ repetitions. During the simulation, the expectation in 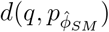 was directly calculated by the numerical integration since the support of *q* is compact. Fig. 13 shows that *B* and 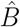 coincides well for various *ϵ*.

**Figure 13:**
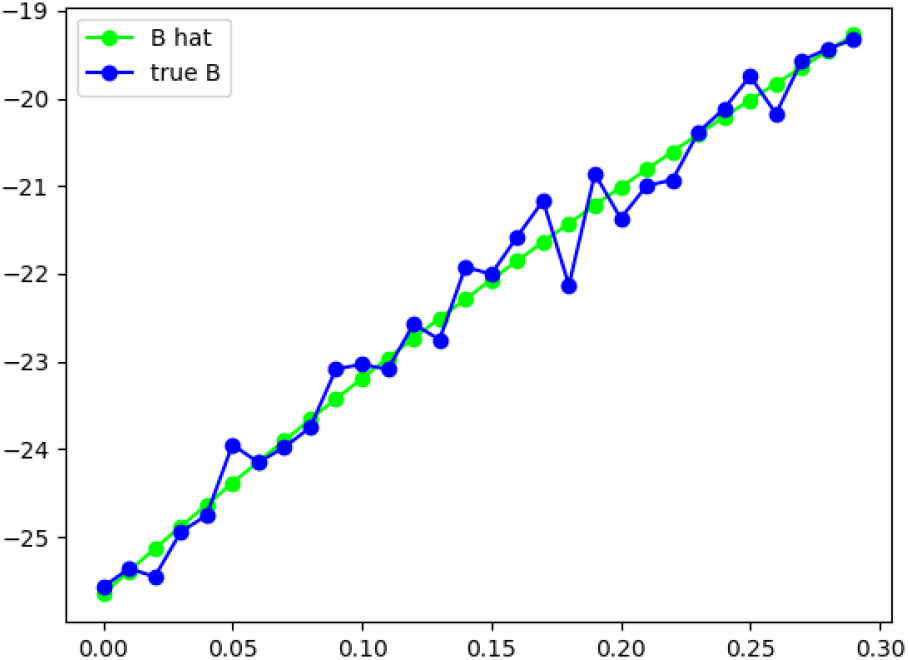
Comparison of *B* and 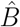 by the Monte Carlo simulation. The axes of the figure are *B* (or 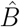) versus *ε*.

### S4 Appendix. Permutation test on synthetic dataset

We have additionally conducted a permutation test on a recovered graph in a simulation experiment using 19-dimensional torus graph, where the settings follow those presented in Section 3.1.

Under the null hypothesis that the *d* variables are mutually independent, we independently permuted the *N* rows within each column — preserving each variable’s marginal distribution while breaking cross-variable alignment — and refit the same graphical model to each permuted dataset. As shown in Fig. 14, the original graph contains significantly more edges than its permuted counterparts.

**Figure 14:**
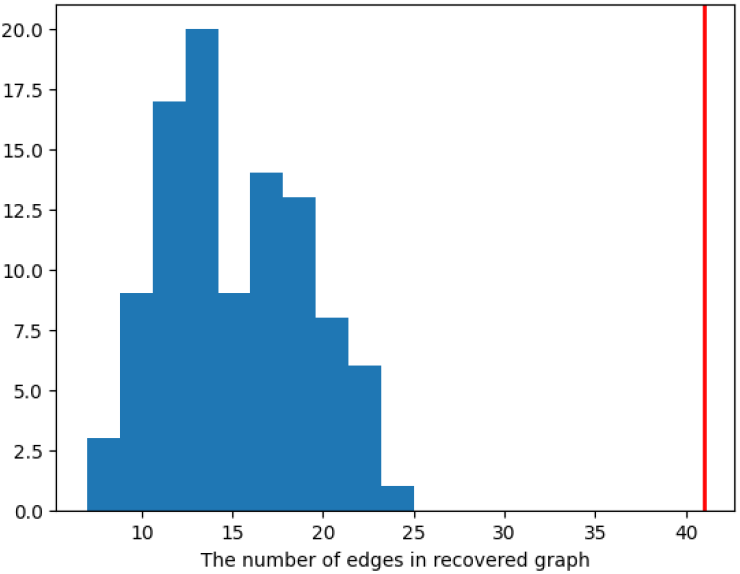
Permutation test on the number of edges in recovered graphs. The red line indicates the results for the original synthetic dataset before permutation.

### S5 Appendix. Comparison between non-sparse and sparse torus graph

To compare results obtained by non-sparse torus graph (Klein et al., 2020) and sparse torus graph (ours) in simulation experiments, we employ an Erdös-Rényi random graph *G*(19, 0.1) as the ground truth network and used 10000 synthetic samples as input. As illustrated in Fig. 15, the non-sparse torus graph is sensitive to thresholding, while the model selection is done automatically based on SMIC in our proposed method. Tab. 11 shows the significant difference in estimated modularity.

**Table 11:** Comparison of estimated modularity obtained with the sparse versus non-sparse torus graph. For the non-sparse torus graph, modularity is evaluated according to the weighted-graph definition.

| Ground Truth | Non-sparse Torus Graph | Ours (elbow) | Ours (minimum SMIC) |
| --- | --- | --- | --- |
| 0.466 | 0.290 | 0.466 | 0.458 |

**Figure 15:**
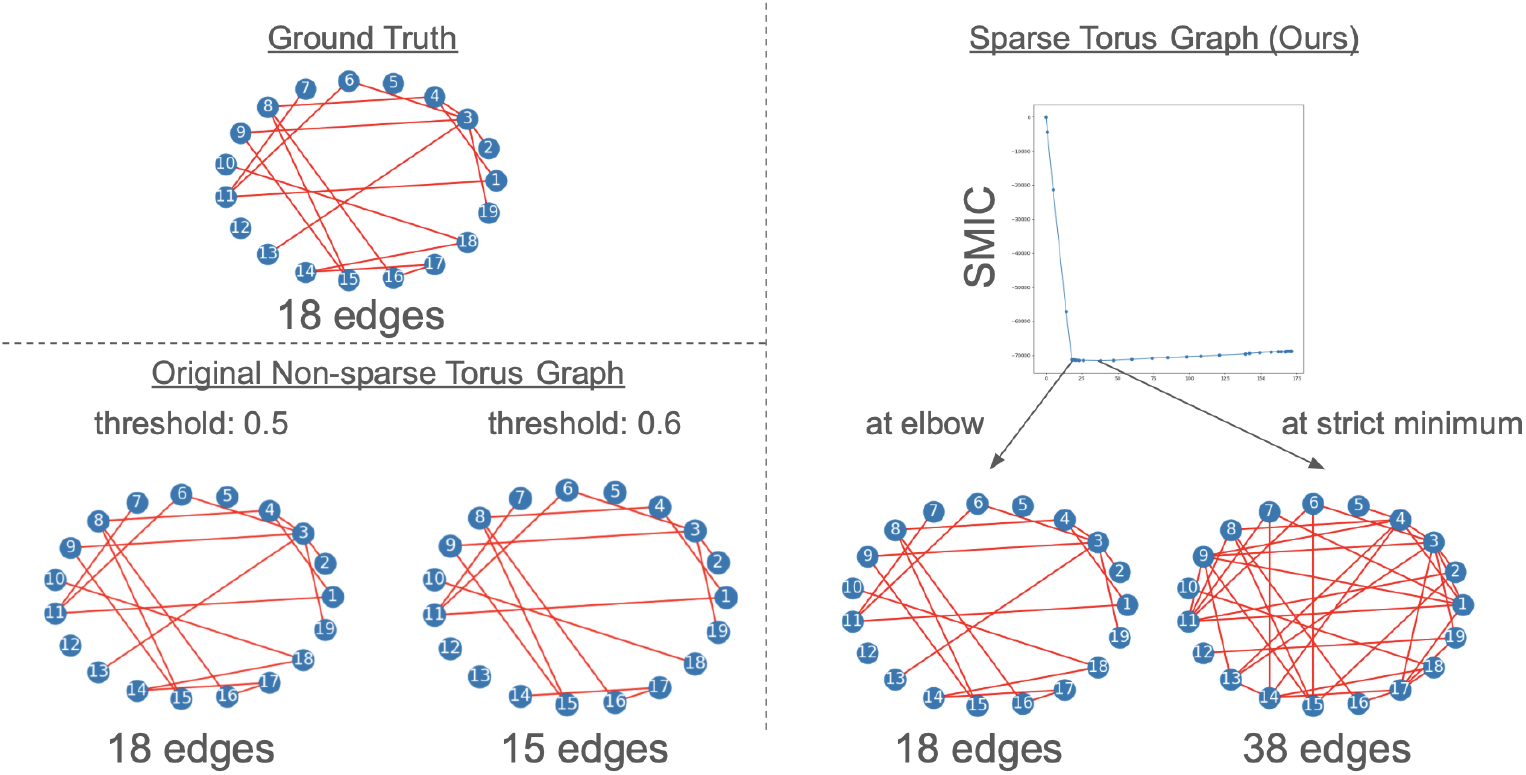
Comparison of results obtained with the non-sparse versus sparse torus graph. The former is sensitive to thresholding.

### S6 Appendix. Full results for human EEG analysis

Full results of the human EEG analysis are shown in Tab.12. Based on these results, the shifts in the recovered graph structures between participant states for the *α, β*, and *γ* bands of human EEG are illustrated in Fig.16, Fig.17, and Fig.18, respectively. Blue lines indicate the 7 patients in the drowsy group, while red lines indicate the 13 patients in the responsive group.

**Table 12:**
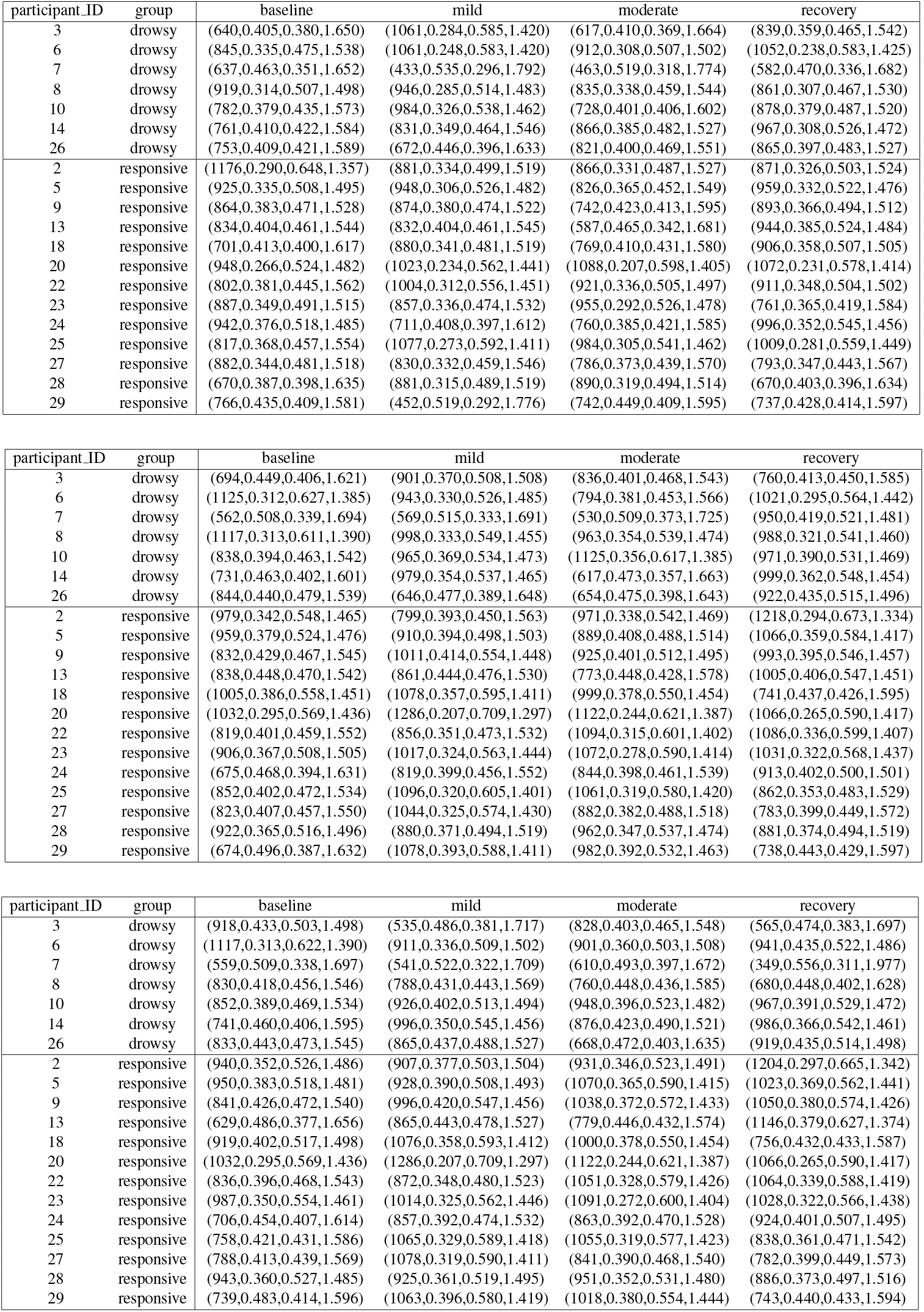
The changes in the number of edges, modularity, average clustering coefficient, and average shortest path length for the recovered graph structures in each state. The first, second, and third table corresponds to *α, β*, and *γ* band, respectively.

**Figure 16:**
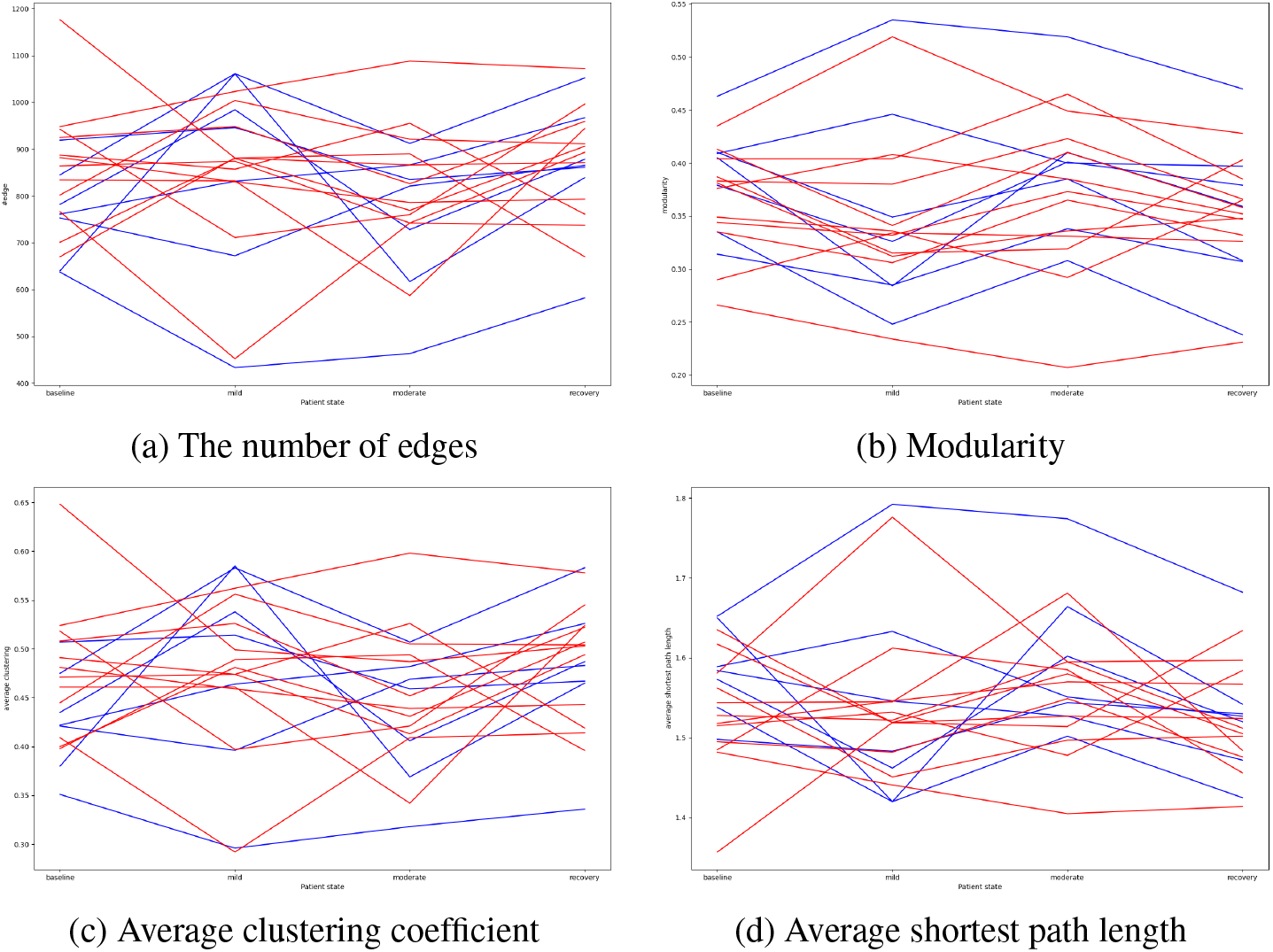
The shift in the recovered graph structures between participant states for *α* band EEGs.

**Figure 17:**
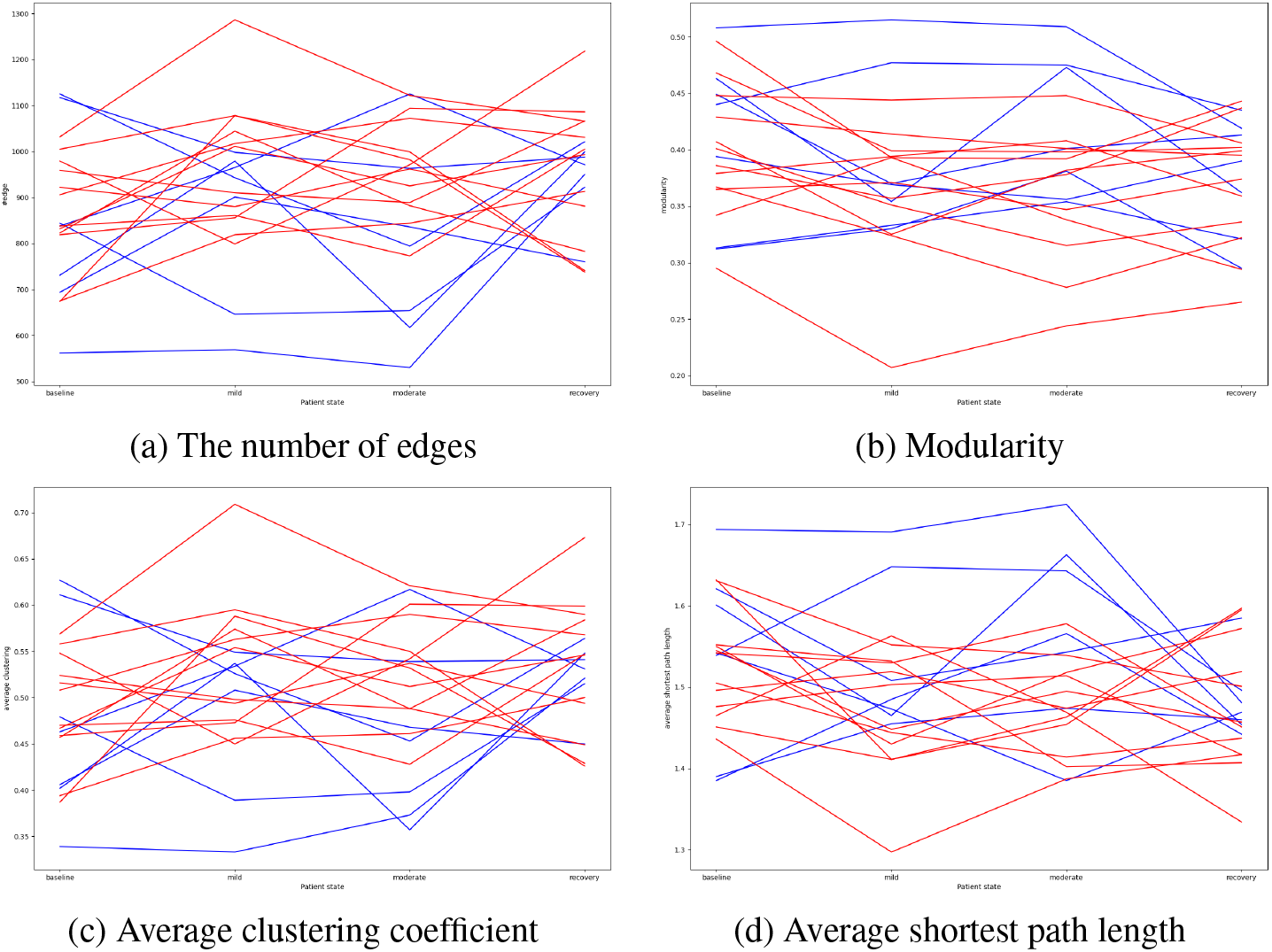
The shift in the recovered graph structures between participant states for *β* band EEGs.

**Figure 18:**
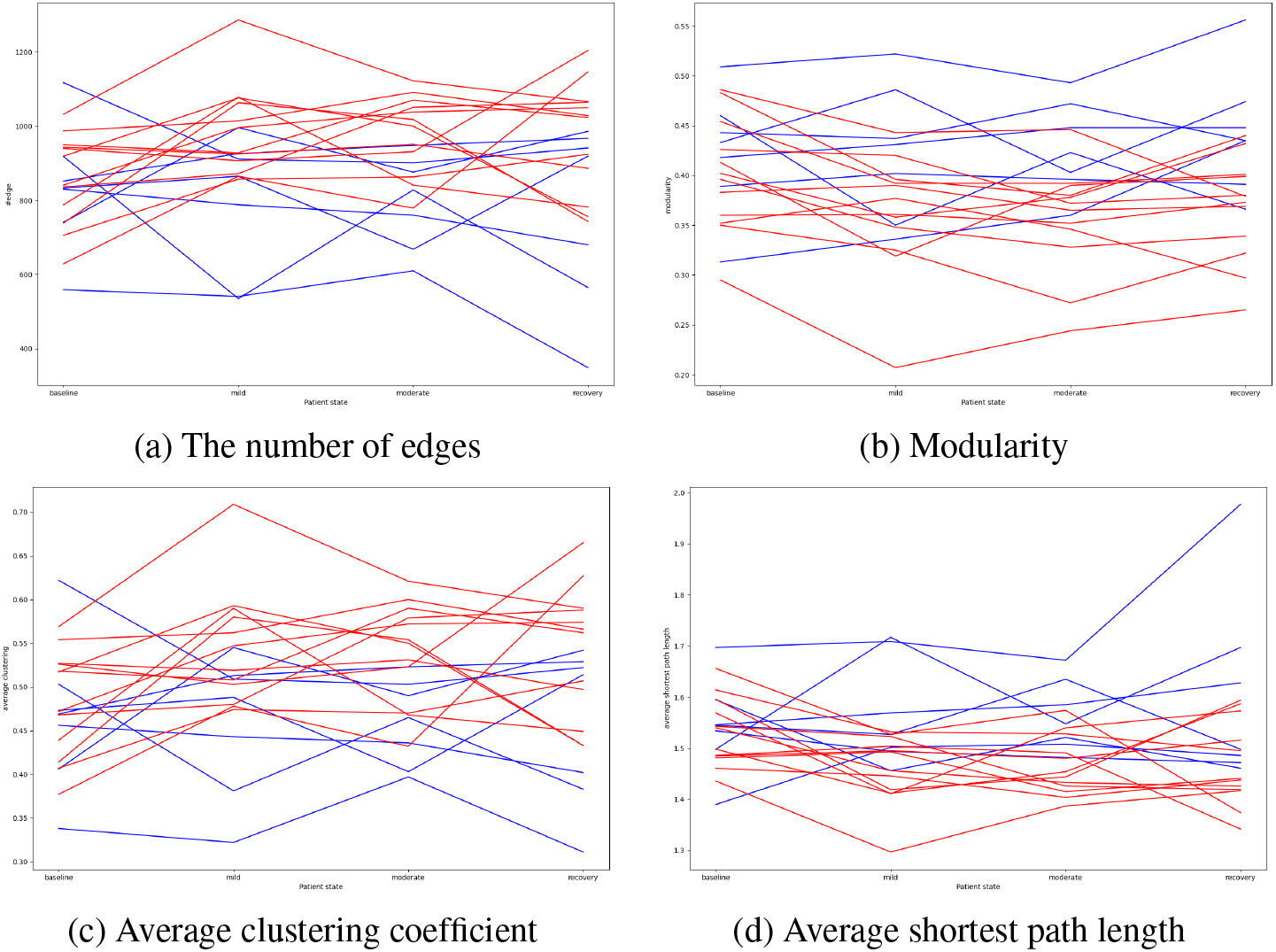
The shift in the recovered graph structures between participant states for *γ* band EEGs.

## Footnotes

1 Note that the term “model” hereafter refers to the statistical model that expresses the data generation probability distribution, which is in contrast to mathematical models, which are typically described using differential equations.

2 Klein et al. (2020) has also mentioned adding a group *l*_1_ penalty in their paper, however practices were not presented.

3 The model with the minimum information criteria is generally considered to fit the data best while suppressing the model complexity. We do not argue the validity of the information criteria itself in this paper.

